# Membrane insertion mechanism of the caveolae coat protein Cavin1

**DOI:** 10.1101/2021.03.23.436578

**Authors:** Kang-cheng Liu, Hudson Pace, Elin Larsson, Shakhawath Hossain, Aleksei Kabedev, Ankita Shukla, Vanessa Jerschabek, Jagan Mohan, Christel A. S. Bergström, Marta Bally, Christian Schwieger, Madlen Hubert, Richard Lundmark

## Abstract

Caveolae are small plasma membrane invaginations, important for control of membrane tension, signaling cascades and lipid sorting. The caveolae coat protein Cavin1 is essential for shaping such high curvature membrane structures. Yet, a mechanistic understanding of how Cavin1 assembles at the membrane interface is lacking. Here, we used model membranes combined with biophysical dissection and computational modelling to show that Cavin1 inserts into membranes. We establish that initial PI(4,5)P_2_-dependent membrane adsorption of the trimeric helical region 1 (HR1) of Cavin1 mediates the subsequent partial separation and membrane insertion of the individual helices. Insertion kinetics of the HR1 is further enhanced by the presence of flanking negatively charged disordered regions, which was found important for the co-assembly of Cavin1 with Caveolin1 in living cells. We propose that this intricate mechanism potentiates membrane curvature generation and facilitates dynamic rounds of assembly and disassembly of Cavin1 at the membrane.

**Significance statement:** Caveolae are cholesterol enriched membrane invaginations coupled to severe muscle and lipid disorders. Their formation is dependent on assembly of the protein Cavin1 at the lipid membrane interface driving membrane curvature. In this work, we dissect the mechanism for how Cavin1 binds and inserts into membranes using a combination of biochemical and biophysical characterization as well as computational modelling. The proposed model for membrane assembly potentiates dynamic switching between shielded and exposed hydrophobic helices used for membrane insertion and clarifies how Cavin1 can drive membrane curvature and the formation of caveolae.

## Introduction

The typical small bulb shaped invaginations of the plasma membrane termed caveolae are found in most vertebrate cells. They are highly abundant in adipocytes, muscle and endothelial cells and are important for various physiological processes like regulation of membrane tension, lipid metabolism and cellular signaling (1, 2). Lack or dysfunction of caveolae are connected to severe human diseases such as muscular dystrophy, cardiomyopathy and lipodystrophy. Caveolae formation is dependent on the membrane lipid composition and the coat components Caveolin1 (CAV1) and Cavin1 (3). Caveolae are enriched in cholesterol and sphingolipids (1, 2), which do not only accumulate in caveolae, but are actively sequestered (4). The negatively charged lipids phosphatidylserine (PS) and phosphatidylinositol (4,5) bisphosphate (PI(4,5)P_2_) are also enriched in caveolae (5). Lipid mapping in cells showed that both CAV1 and Cavin1 recruit specific lipid species to caveolae, hereby acting synergistically to generate the unique lipid nano-environment of caveolae (6, 7). CAV1 and Cavin1 are universal structural elements, and knock out of either of these proteins leads to loss of caveolae (1, 2). Electron microscopy studies on caveolae have revealed a striated protein coat lining, which is believed to comprise CAV1 and the Cavin proteins (8, 9). CAV1 belongs to a family of integral membrane proteins (CAV1-3), where both the N- and C-terminal protrude into the cytoplasm. CAV1 has been shown to form high order 8S oligomers in membranes following cholesterol binding (10). Cavin1 belongs to a family comprised of four different proteins (Cavin1-4), which exhibit tissue specific expression patterns (3). The Cavin proteins are thought to assemble with the CAV1 8S complexes to form 60S and 80S complexes building up the caveolae coat (11). Importantly, Cavin1 is required for membrane invagination of caveolae (12). Cryo-electron microscopy studies of such complexes proposed an architecture composed of an inner cage of polygonal units of Caveolins, and an outer Cavin coat (13, 14). The models propose that Cavin arranges into a web-like architecture composed of an interbranched trimeric complex (13) or alternatively that the Cavins are stacked in rod-like trimers (14). However, it is still not understood how the unique striped or spiral pattern of the caveolae coat is assembled and what intermolecular forces join the molecular components together.

The Cavin proteins share a common pattern in their domain structure, containing negatively charged disordered regions (DRs) interspersed with positively charged helical regions (HRs) (Figure 1A). The crystal structures of the HR1 (4QKV and 4QKW) revealed an extended α-helical trimeric coiled-coil structure (15). The HR1 domain has been shown to mediate trimeric homo-oligomerization of Cavin1, and formation of hetero-complexes with either Cavin2 or Cavin3 in solution (15, 16). The HR2 is also thought to build up a trimeric coiled coil, but this structural arrangement is dependent on the HR1. *In vitro* studies have shown that Cavin1 binds both PI(4,5)P_2_ and PS (15, 17). The positively charged amino acids (Lys115, Arg117, Lys118, Lys124, Arg127) in the HR1 domain mediate specific binding to PI(4,5)P_2_ (15), whereas a repeated sequence of 11 amino acids of the HR2 domain, identified as undecad repeat (UC1), was shown to bind PS (17). Furthermore, Cavin1 has been shown to generate membrane curvature *in vitro* (15). Both HR and DR were required for this, and it was proposed that Cavin1 drives membrane curvature by molecular crowding via weak electrostatic interactions between the DR and HR regions (18). Interestingly, the assembly of both CAV1 and Cavin1 was found to be dependent on the acyl chain composition of PS, suggesting that Cavin1 might also interact with the hydrophobic region of the membrane (6). Membrane insertion of Cavin1 could contribute to membrane curvature generation and the formation of caveolae. Yet, based on the current structural understanding, it is not clear how Cavin1 orients and assembles at the membrane interphase.

**Fig. 1.**
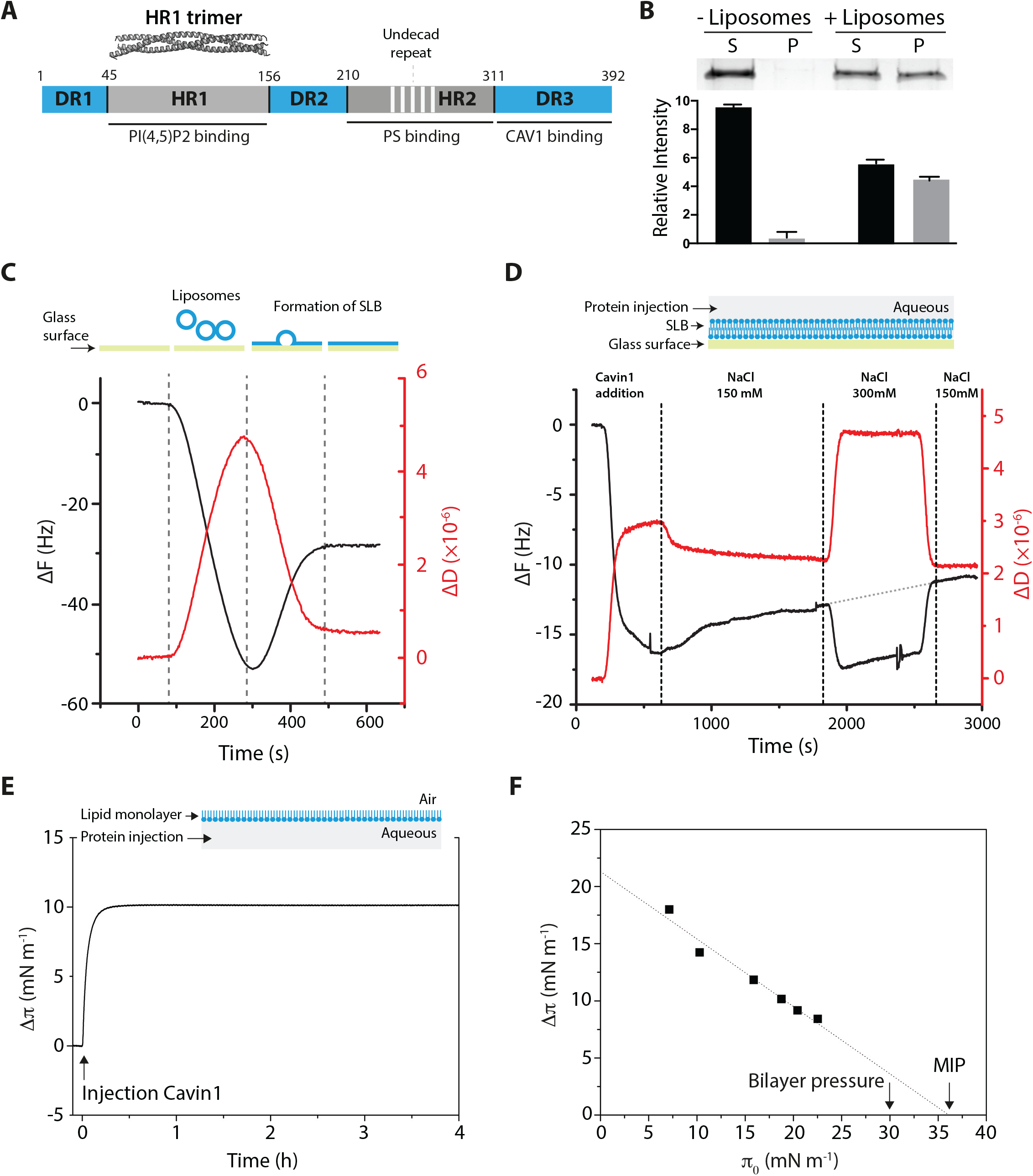
Cavin1 binding and insertion into model lipid membranes. **(A)** Scheme of the domain-structure of Cavin1 with DR and HR. White stripes mark undecad repeats. The crystal structure of HR1 (4QKV) is displayed on top. Regions involved in binding to PI(4,5)P_2_, PS ad CAV1 are indicated. **(B)** Liposome co-sedimentation of Cavin1. Cavin1 was incubated with or without DOPC:DOPE:PI(4,5)P_2_ liposomes, centrifuged, and supernatant (S) and pellet (P) fractions were analyzed by SDS-PAGE. Band intensities were quantified and data is shown as mean ± SEM (n=3). **(C)** Top; scheme of SLB formation. Bottom; QCM-D measurement showing shift in frequency (ΔF) (black line) and dissipation (ΔD) (red line) upon SLB formation. **(D)** Top; illustration of QCM-D set up. Bottom; QCM-D monitoring of Cavin1 adsorption to a SLB. The responses in ΔF and ΔD are corresponding to Cavin1 injection and buffer rinses as indicated. Grey dotted line shows extrapolation of protein desorption from first rinse (150 mM NaCl). **(E)** Top; scheme of monolayer protein adsorption experiments. Bottom; Cavin1 adsorption to DOPC:DOPE:PI(4,5)P_2_ monolayers. Cavin1 was injected underneath the film at π_0_ = 20 mN m^-1^ and Δπ recorded over time. **(F)** Cavin1 adsorption to lipid monolayers was measured at different π_0_. The MIP value was determined by extrapolation of the Δπ/π_0_ plot to the x-axis.

In this work, we address the detailed mechanism by which Cavin1 binds and assembles at the lipid interface using model membranes in combination with a variety of biophysical techniques. We found that Cavin1 inserted into the membrane via the HR1 domain in a PI(4,5)P_2_-mediated process. Membrane insertion involved partial separation of the helices in the HR1 regions in a process aided by the DR domains.

## Results

### Cavin1 inserts into the membrane providing stable membrane association

To characterize the mechanism of membrane-driven assembly of Cavin1, we purified full-length Cavin1 from mammalian human embryonic kidney (HEK) cells (Fig. S1A and B). Analysis of binding to liposomes composed of DOPC:DOPE:PI(4,5)P_2_ (55:45:5 mol%) using a co-sedimentation assay showed that Cavin1 bound to membranes with relatively high positive curvature (Fig. 1B). To further address the membrane association in a system independent of membrane curvature, and where binding over time could be quantified, we generated flat supported lipid bilayers (SLBs) composed of POPC:PI(4,5)P_2_ (95:5 mol%) on a glass surface. Quartz crystal microbalance with dissipation monitoring (QCM-D) has been used extensively to monitor SLB formation (19), as it provides a distinct signature of changes in the adsorbed mass (frequency, ΔF) and stiffness (dissipation, ΔD) of the surface (Fig. 1C). Addition of purified Cavin1 to the SLB resulted in a steep decrease in frequency and a concerted increase in dissipation, representing adsorption of the protein to the SLB (Fig. 1D). When the system was rinsed with buffer (150 mM NaCl), we noticed a slow but steady release of Cavin1 from the SLB surface, indicating that the binding is at least partially reversible. We next tested if Cavin1 was adsorbed to the SLB through electrostatic interactions by treating the system with increased salt concentration. The change to buffer containing 300 mM NaCl caused a sudden shift in both the frequency and dissipation due to the difference in densities between the two buffers, but was reversed upon switching back to the initial buffer. We found that higher concentration of salt did not cause additional protein desorption from the SLB beyond the slow rate observed prior to the high salt treatment (Fig. 1D, grey dotted line). This suggested that, once bound to the membrane, Cavin1 was not solely interacting with the SLB through electrostatic interactions. In contrast, increased salt concentration during adsorption of Cavin1 to the SLBs led to a ∼30% reduction in the amount of protein that bound (Fig S1C); thereby indicating that electrostatic interactions are important for the initial recruitment of Cavin1 to the membrane.

To determine if Cavin1 inserts into membranes we used a Langmuir trough. This technique monitors the adsorption of a protein to a lipid monolayer suspended at an air/water interface through changes in the lateral pressure of the monolayer, thereby indicating the degree with which a protein is inserting into the monolayer (20). The surface pressure (π) is directly related to the lateral cohesion of molecules and initial surface pressure values (π_0_) are related to the lateral packing density of the lipids before protein interactions. We prepared lipid films of DOPC:DOPE:PI(4,5)P_2_ (55:45:5 mol%) on a buffer surface and injected Cavin1 underneath the lipid monolayer into the subphase (Fig. 1E). The rapid increase in surface pressure seen following protein injection indicated that Cavin1 instantly adsorbed to and inserted into the lipid monolayer (Fig. 1E). To get further insights into binding mechanisms, we measured Cavin1 adsorption at various initial surface pressures (π_0_) and monitored surface pressure variation (Δπ) induced by protein/lipid interaction (Fig. S1D). Linear regressions of Δπ = f(π_0_) provides a synergy factor (*a*) that corresponds to the slope +1 and describes the protein affinity for the lipid monolayer (Fig. 1F). The positive synergy factor value of *a* = 0.41 for Cavin1 indicated the existence of strong protein/lipid interactions (21, 22). This is further supported by a high maximum insertion pressure (MIP) value of 36.1 ± 1.9 mN m^-1^ obtained by extrapolation of the adsorption data to the x-axis (Fig. 1F). Proteins with MIP values above the estimated monolayer/bilayer equivalence pressure (∼30 mN m^-1^) are considered to be well incorporated into the lipid layer (23). Our data indicated a high extent of Cavin1 membrane insertion, likely due to a combination of both electrostatic and hydrophobic forces.

### The N-terminal region of Cavin1 adsorbs and inserts into membranes in a PI(4,5)P_2_-dependent manner

To identify the region required for membrane insertion of Cavin1, we expressed and purified the N-terminal part (1-190) and the C-terminal part (191-392) of Cavin1 as recombinant proteins from bacteria (Fig. S2A). By performing a co-sedimentation assay we found that the truncated Cavin1 (1-190) variant bound membranes equally well as the full-length protein (Fig. 2A and Fig. 1B, respectively). However, as previously shown, the C-terminal region (191-392) was unable to bind to liposomes composed of DOPC:DOPE:PI(4,5)P_2_ (17). To test whether the 1-190 region of Cavin1 was sufficient for membrane insertion, we performed adsorption experiments using lipid monolayers. The data revealed a remarkable increase in surface pressure following injection of Cavin1 (1-190) underneath a DOPC:DOPE:PI(4,5)P_2_ film, showing that this part of the protein does indeed insert (Fig. 2B). While the synergy factor of *a* = 0.43 was similar to that of full-length protein (*a* = 0.41), the MIP was determined to be 49.6 ± 2.2 mN m^-1^ in comparison to 36.1 ± 1.9 mN m^-1^ for Cavin1 FL, confirming that Cavin1 (1-190) has a high affinity for the lipid interface and is strongly incorporated into the monolayer (Fig. 2C and Fig. S2B).

**Fig. 2.**
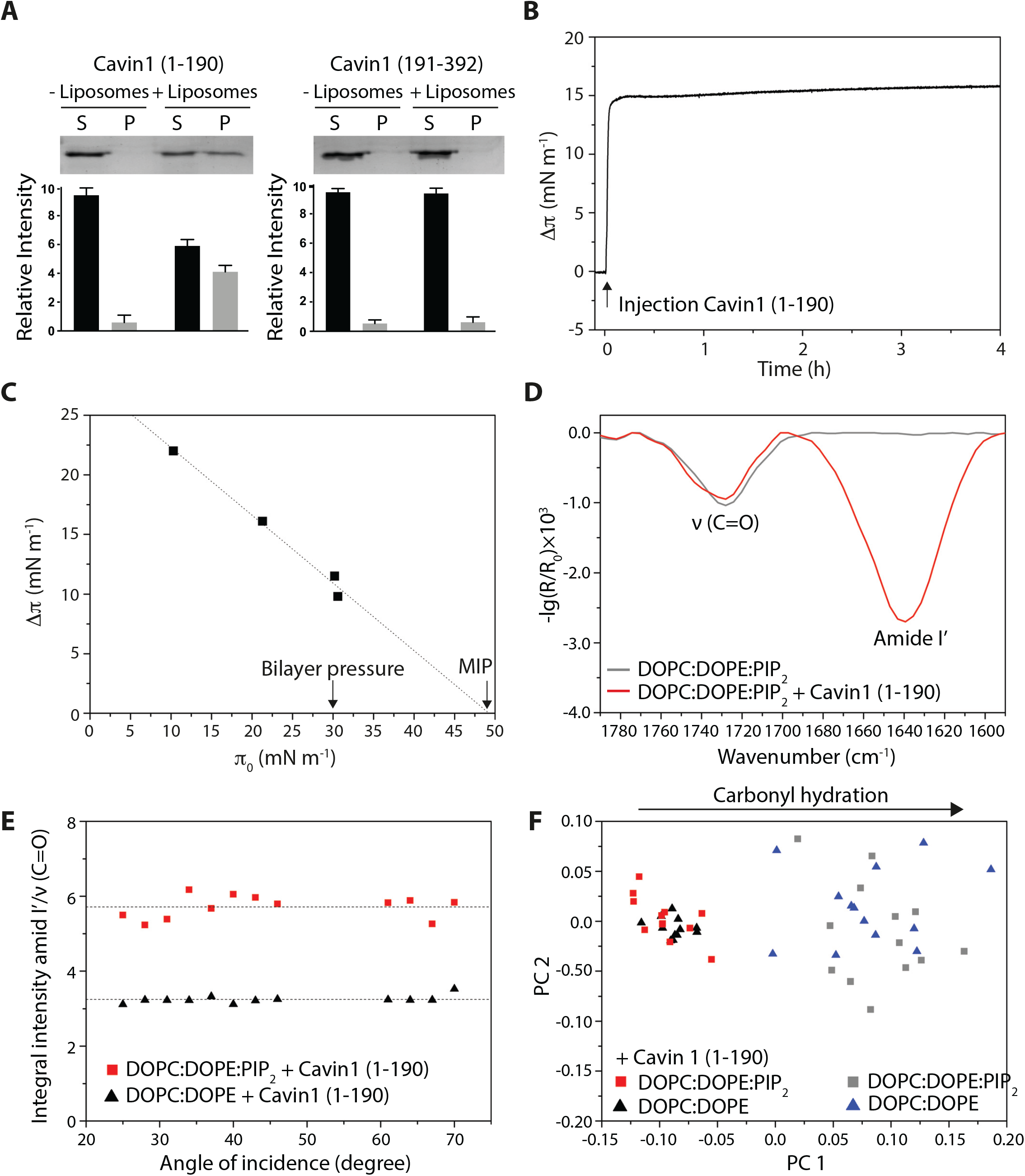
The amount of Cavin1 (1-190) inserted into membranes is PI(4,5)P_2_ dependent. **(A)** Liposome co-sedimentation assay. Cavin1 (1-190) or Cavin1 (191-392) were incubated with or without liposomes, centrifuged, and supernatant (S) and pellet (P) fractions were analyzed by SDS-PAGE. Band intensities were quantified and data is shown as mean ± SEM (n=3). **(B)** Cavin1 (1-190) adsorption to DOPC:DOPE:PI(4,5)P_2_ monolayers. Cavin1 (1-190) was injected at π_0_ = 20 mN m^-1^ and Δπ recorded over time. **(C)** Cavin1 (1-190) adsorption to lipid monolayers was measured at different π_0_. MIP was determined by extrapolation of Δπ/π_0_ plot to x-axis. **(D)** IRRA spectra (1790–1590 cm^-1^) of DOPC:DOPE:PI(4,5)P_2_ at π_0_ = 20 mN m^-1^. The C=O vibrational band (∼1730 cm^-1^) originates from lipid ester groups. The amide I’ band (∼1640 cm^-1^) indicates Cavin1 (1-190) adsorption after injection into the subphase. Spectra were acquired with p-polarized light at an angle of incidence of 40°. **(E)** Amount of protein adsorbed to the monolayer. Ratios of integral intensity of amide I’ and C=O bands are shown as a function of the angles of incidence for indicated lipid monolayers. Dotted lines display the mean of each data set. **(F)** PCA in the C=O vibrational region based on IRRA spectra of DOPC:DOPE:PI(4,5)P_2_ and DOPC:DOPE monolayers before and after adsorption of Cavin1 (1-190). PC1 represents the extent of carbonyl group hydration (76% of total variance), while PC2 (10%) showed no additional systematic changes.

To investigate the effect of protein insertion on membrane organization, we used infrared reflection-absorption spectroscopy (IRRAS). IRRA spectra of the pure lipid films consisting of either DOPC:DOPE:PI(4,5)P_2_ (Fig. 2D) or DOPC:DOPE (Fig. S2C) displayed characteristic C=O vibrational bands at ∼1730 cm^-1^ originating from the ester group of the lipids. After injection of Cavin1 (1-190), the presence of the protein at the air/buffer interface was indicated by the amide I’ band with a maximum at ∼1640 cm^-1^ (Fig. 2D and Fig. S2C). The intensity of amide I’, which correlates to the amount of adsorbed protein, was increased when PI(4,5)P_2_ was present in the lipid layer. To assess the amount of inserted protein per lipid molecule, the ratios of integral intensities of amide I’/ν(C=O) of various IRRA spectra recorded at different angles of incidence were calculated for both lipid monolayers after Cavin1 (1-190) adsorption (Fig. 2E). The data showed significantly higher protein/lipid ratios for DOPC:DOPE:PI(4,5)P_2_ films (amide I’/ν(C=O) = 5.71 *vs*. 3.24, \**p* < 0.05) indicating a strong correlation between the presence of PI(4,5)P_2_ and the amount of adsorbed Cavin1 (1-190). To study how insertion of Cavin1 (1-190) affects the lipid monolayer, we used the frequency of the carbonyl vibration, which is sensitive to H-bond formation and thus provides information on the lipid hydration at the hydrophilic/hydrophobic interface (24). Principal component analysis (PCA) of a large number of recorded spectra revealed that the differences are indeed due to different extents of hydration (Fig. S2D). Pure lipid monolayers displayed more hydrated carbonyls and the binding of Cavin1 (1-190) resulted in less hydrated carbonyl groups (Fig. 2F). Moreover, IRRA spectra of pure lipid films at the surface pressure of injection (22 mN m^-1^) and after protein adsorption (36 mN m^-1^) (Fig. S2F) were similar. This indicated that lipid dehydration is not an effect of increasing surface pressure following Cavin1 injection but can clearly be assigned to insertion of Cavin1 (1-190) into the lipid head group region and the concomitant replacement of hydration water.

### The HR1 is membrane bound in a slightly inclined angle in the presence of the DR1 and DR2

To further address how membrane binding and insertion of Cavin1 (1-190) would affect the protein structure, we used far-UV circular dichroism (CD) spectroscopy. We found that Cavin1 (1-190) exhibited a predominantly α-helical CD profile with typical minima at 208 and 222 nm at both 300 mM and 150 mM NaCl (Fig. 3A and B). However, following preincubation with liposomes we noticed a dramatic change in the CD spectra at 150 mM NaCl, but not at 300 mM NaCl (Fig. 3A and B). This involved a significant increase of the ellipticity at 222 nm in relation to 208 nm (Fig. 3B), which is indicative of an increased hydrophobicity of the helix environment. This could derive from either membrane insertion or oligomerization, as both would protect the helical surfaces from the solvent. To further monitor how the HR1 domain, in combination with the DR1 and DR2 domains, would insert and orient at the membrane, we used IRRAS to get direct information on the secondary structure of the adsorbed protein and its orientation (20). The secondary structure of the protein is encoded in the position of the amid I’ band, whereas the orientation influences its intensity. Knowledge of the protein structure is essential for appropriate data analysis and determination of the orientation at the lipid monolayer. Since the crystal structure of the HR1 (45-155) is known (15) and both DR1 (1-43) and DR2 (156-190) lack a secondary structure, we could use this technique to understand the membrane association of Cavin1 (1-190) in more detail. The asymmetric amid I’ band shape in the experimental spectra indeed indicated that in addition to helical structures, further components such as unordered structural elements were present (Fig. 3C). Moreover, the positions of the helical components are typical for a coiled coil structure (25). This is in agreement with the combination of HR and DR domains (15). To predict orientation of Cavin1 (1-190) when adsorbed to a DOPC:DOPE:PI(4,5)P_2_ monolayer, experimental and simulated IRRA spectra recorded at various angles of incidence with p- or s-polarized IR-light were compared (Fig. 3C). The best fitting band simulation yielded an average inclination angle of *γ* = 22.5 ± 2.5° for the helical components of Cavin1 (1-190) with respect to the lipid monolayers (Fig. 3C). However, upon fitting simulated to experimental spectra at all theoretically possible inclination angles, we found that the minimum is rather shallow and spectral fits are reasonable for average helix inclination angles between 0° (parallel to the lipid layer, Fig. S3A) and 30° (slightly inclined, Fig. 3D). Conversely, at higher inclination angels (γ > 30°) no acceptable spectral fit could be obtained (Fig. S3B-C). It should be noted that an inclination angle of γ = 0° could only be obtained if the HR1 helix bundle dissociates and all three helices adsorb individually and horizontally to the interface. If HR1 were to adsorb as an intact trimer, the smallest possible helix inclination angle, would be γ = 12° due to the intrinsic helix orientations within the bundle. The minimum found in the IRRA spectra fit corresponds to a HR1 bundle inclination of 20 ± 3° (Fig 3D). Our results implied that the HR1 domain of Cavin1 (1-190) adsorbs with a slight inclination to the interface, where average inclination angles of 17–23° were most probable (Fig. 3D).

**Fig. 3.**
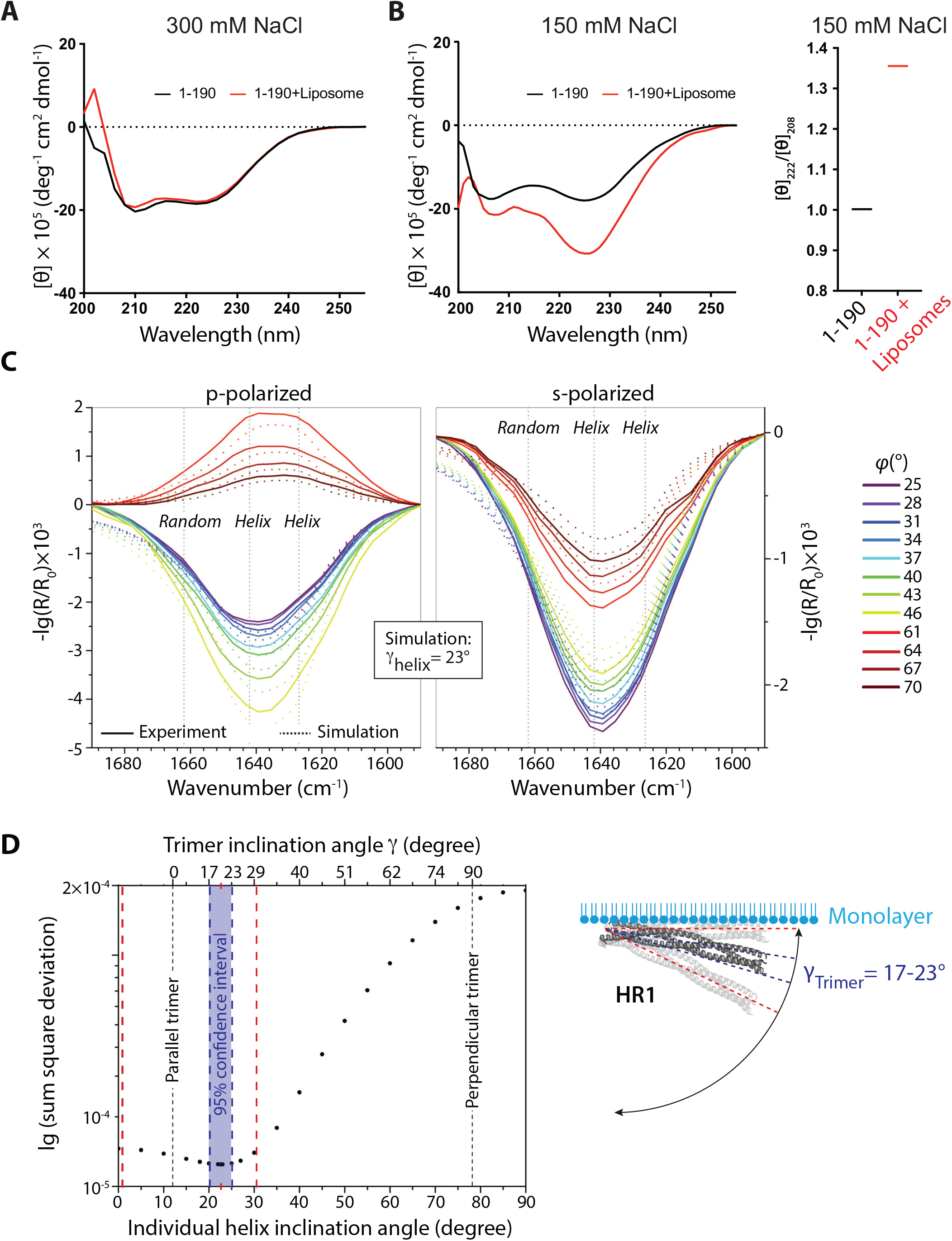
Adsorption of Cavin1 (1-190) to membranes involves inclination of the helices in relation to the membrane. **(A-B)** Far-UV CD spectra (195–255 nm) of Cavin1 (1-190) with or without liposomes in 300 mM **(A)** or 150 mM **(B)** NaCl buffer. In (B), the ratio between 222 and 208 nm ellipticities ([θ_222_]/[θ_208_]) in the absence and presence of liposomes is plotted to the right. **(C)** Experimental and best fitting simulated IRRA spectra of Cavin1 (1-190) adsorbed to a DOPC/DOPE/PI(4,5)P_2_ monolayer, at various angles of incidence *φ* and polarizations. Vertical dotted lines indicate the band components used for simulation. The best fit was achieved with a helix inclination angle of γ = 22.5 ± 2.5° with respect to the lipid interface. **(D)** The quality of the fit was assessed as sum square deviation between experimental and simulated spectra at various γ. The minimum is indicated by a red x-axis tick. The 95% confidence interval of the minimum is shown in blue. Red vertical lines mark a slightly larger range of plausible HR1 orientations considering uncertainties in experiment, model assumptions and data treatment. The upper x-axis shows the inclination angles of the complete HR1 trimer that correspond to the individual helices’ inclination angles. Note that their relation is not linear. Right panel in (D) shows scheme of HR1 trimer orientation at lipid monolayer. The most probable orientation is shown in dark gray and outlined by blue dotted lines. Red dotted lines indicate the range of inclination angles that plausibly explain the measured spectra.

### The DR1 and DR2 influence the membrane binding properties of the HR1

To further examine the mechanism for how the N-terminal region (1-190) interacts and inserts into the membrane, we generated and purified different truncated versions of the 1-190 region from *E. coli* (Fig. 4A and Fig. S4A). Using CD spectroscopy, we found that all constructs containing the HR1 exhibited a predominantly α-helical CD profile in 300 mM NaCl buffer with typical minima at 208 and 222 nm (Fig. S4B). However, at 150 mM NaCl concentration the 44-190 construct lost the α-helical CD profile (Fig. S4C). To analyze the oligomeric state of the constructs, we used mass photometry (Fig. 4B). All constructs, where detected as trimers, although the measured molecular weight of all Cavin1 constructs was slightly higher than theoretically expected, likely due to the elongated structure of the HR1 (Fig 4B). Notably, (1-190) was also detected as monomers, suggesting that the DR1 and DR2 might destabilize the HR1. Liposome co-sedimentation analysis showed that (1-190), (44-155), (1-155) and (101-190) bound membranes, suggesting that the 101-155 region of the HR1 is the minimal part of the protein required for adsorption (Fig. 4C). This region includes the positively charged patch of the HR1, previously shown by mutational analysis to convey PI(4,5)P_2_ binding (15). Notably, the (44-155) construct displayed more membrane binding as compared to constructs that contained the DR1 and/or DR2 in addition to the HR1 (Fig. 4D). To further assess this, we assayed the adsorption of Cavin1 (1-190), (1-155), (44-155) and (44-190) to SLBs using QCM-D. Interestingly (1-155), (44-155), and (44-190) demonstrated much faster adsorption kinetics as well as a greater overall amount of protein adsorbed (−34.7±1.2, -36.4±1.5, and -31.0±4.7 Hz, respectively) than was observed for (1-190) (−25.1±3.4 Hz) (Fig. 4E). However, once the systems were rinsed with buffer, much faster desorption kinetics were also observed for (1-155), (44-155), and (44-190) in comparison to (1-190) (Fig. 4E). These data clearly show that the presence of DR1 and DR2 does not only affect how HR1 initially binds to the membrane, but is also important in determining how strongly the protein is retained; which in turn could be tied to either the degree of membrane insertion or protein-protein interactions. In order to better understand the impact of how these different regions lead Cavin1 to interact with the membrane, we calculated the softness of the adsorbed protein layer (ΔD/-ΔF) after rinsing and at equilibrium (Fig. 4F). The constructs without the DR1 formed much stiffer/denser layers than constructs containing the DR1. This observed difference in the rigidity of the protein coated membrane indicates that the DR1 and DR2 greatly influence not only how the binding of HR1 to the membrane, but also affects its final conformation on the surface.

**Fig. 4.**
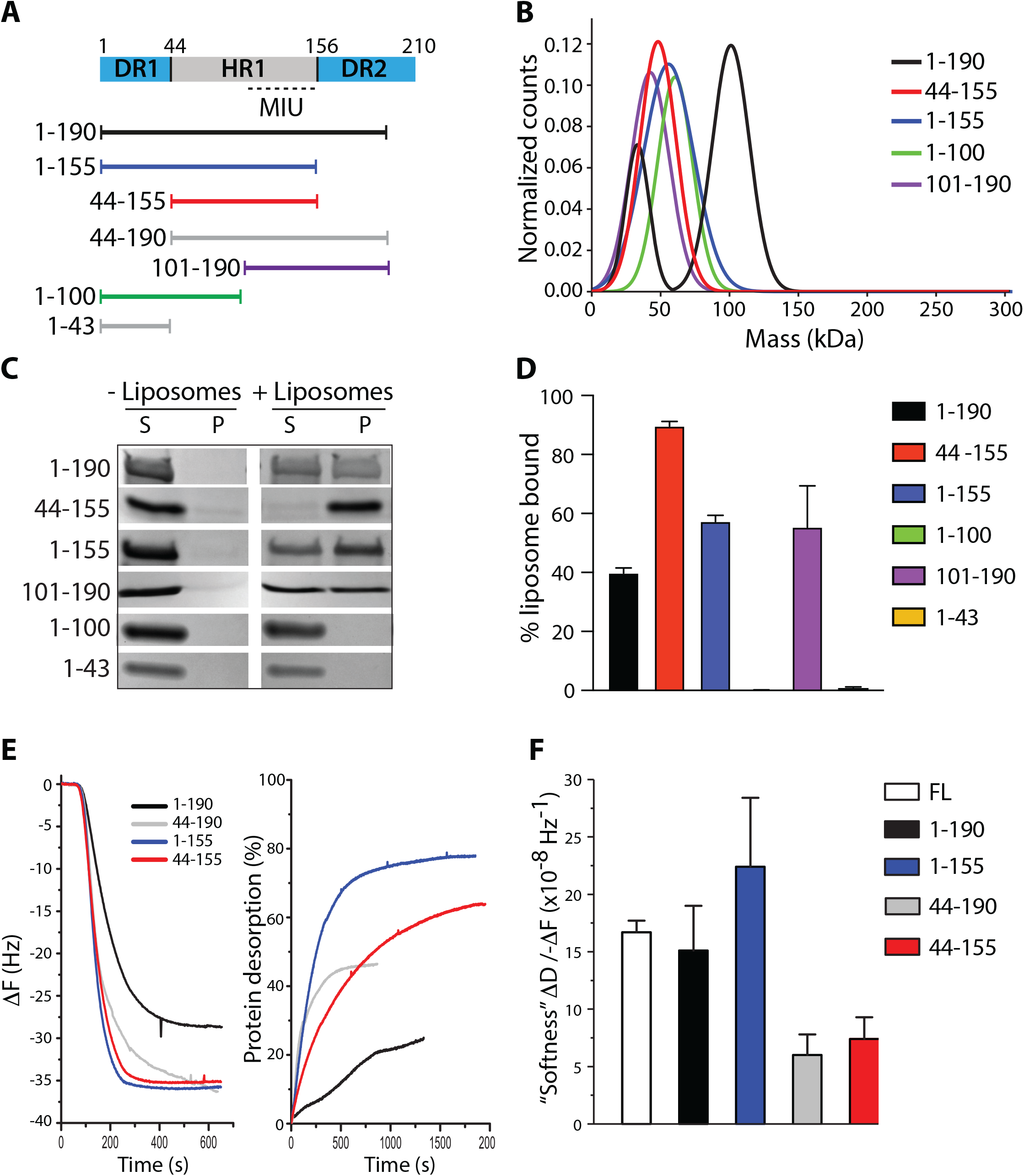
DR1 and DR2 influence the membrane binding properties of the HR1. **(A)** Scheme of the Cavin1 N-terminal domains and purified protein constructs with defined amino acids numbers as indicated. **(B)** Gaussian fit of the molecular weight of Cavin1 constructs as determined using mass photometry. **(C)** Representative liposome co-sedimentation of Cavin1 constructs as indicated. Proteins were incubated with or without DOPC:DOPE:PI(4,5)P_2_ liposomes, centrifuged and supernatant (S) and pellet (P) fractions were analyzed by SDS-PAGE. **(D)** Band intensities of liposome co-sedimentation assay were quantified and data is shown as mean ± SD (n=3). **(E)** Frequency shift (ΔF, left panel) as a result of adsorption of Cavin1 constructs onto SLBs measured by QCM-D. Extent of protein desorption (right panel) following rinse with 150 mM NaCl buffer. **(F)** Softness of the different adsorbed protein layers was calculated as (ΔD/-ΔF) after rinsing and at equilibrium. Data is shown as mean ± SD (n ≥ 2).

### The membrane insertion unit (MIU) of the HR1 is buried into membranes

To study if the different truncated proteins would insert into membranes, their adsorption to monolayers was monitored. Injection of Cavin1 (1-190), (1-155) and (101-190) resulted in an immediate steep increase in surface pressure (Fig. 5A and 5B). These data suggested that the (101-155) residues within the HR1 region, hereinafter referred to as the membrane insertion unit (MIU) (Fig 4A), is responsible for insertion into membranes. Indeed, the MIP of (101-190) was similar to full length Cavin1 (Fig. S5A, 35.5 mN m^-1^ *vs*. 36.1 mN m^-1^, respectively). Interestingly, the (44-155) and (44-190) constructs, which both contain the HR1 but lack the DR1, resulted in a lower surface pressure increase and considerably lower rate of insertion (Fig. 5A). These data suggest the DR1 significantly contribute to the ability of the MIU to insert into the monolayer in the presence of the entire HR1. The DR1 (1-43) alone, or a 1:1 mixture of DR1 (1-44) and HR1 (45-155) domain resulted in a low surface pressure increase (Fig. S5B), showing that a direct linkage between the DR1 and/or DR2 with HR1 is required for efficient insertion.

**Fig. 5.**
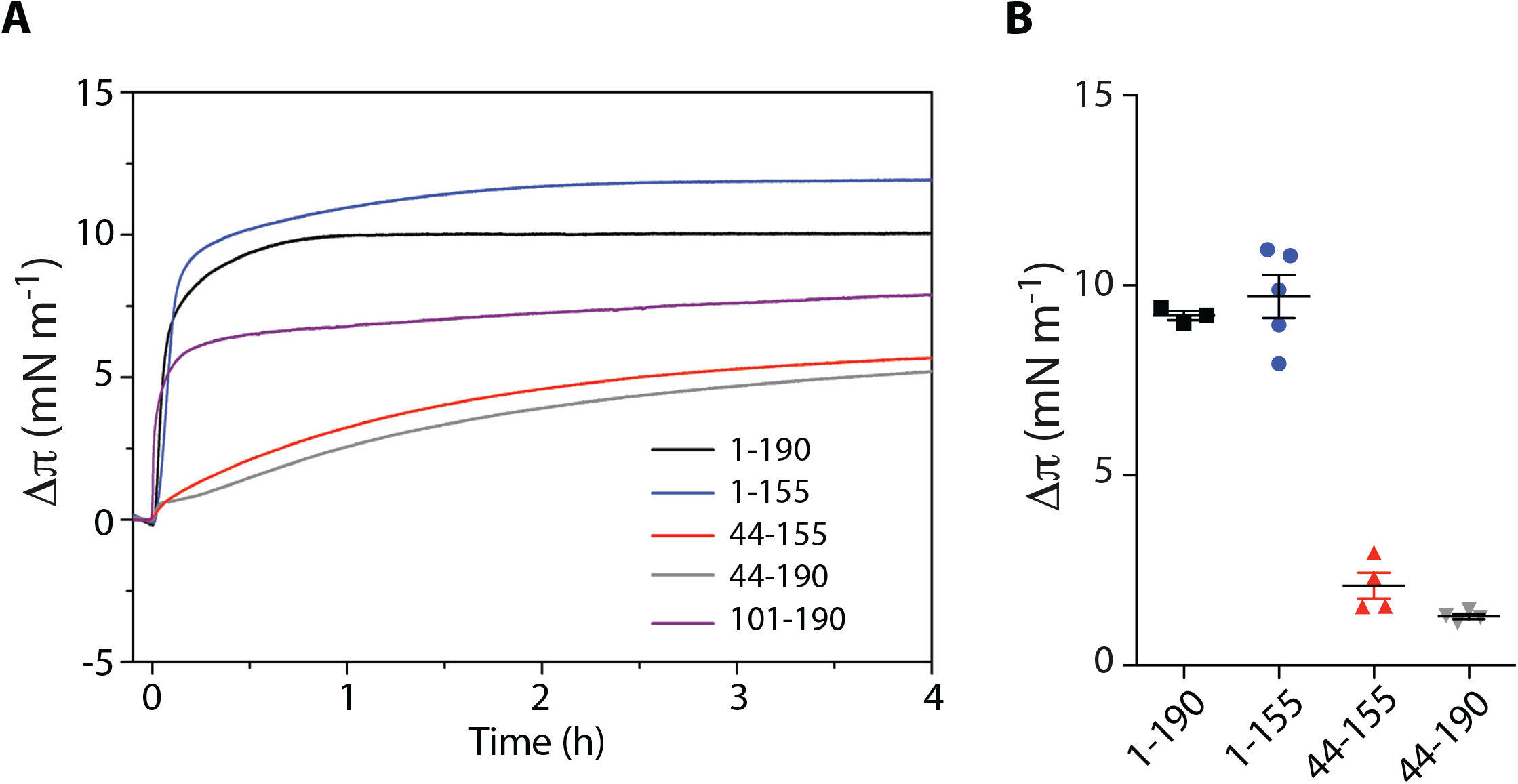
The membrane insertion unit (MIU) of the HR1 is buried into membranes. **(A)** Adsorption of Cavin1 truncates as indicated to DOPC:DOPE:PI(4,5)P_2_ monolayers. Proteins were injected at π_0_ = 20 mN m^-1^ and Δπ recorded over time. Dotted line indicates the time-point for quantification in (B). **(B)** Quantification of Δπ recorded in (A) at t = 20 min. Data is shown as mean ± SEM.

Previous work has shown that removal of the DR1 from full length Cavin1 affected the ability of the protein to efficiently co-assemble with CAV1 in cells (18). We could confirm these results showing that Cavin1-ΔDR1 only partially assembled together with CAV1 at the plasma membrane, but that the majority of CAV1 was present in larger internal membranous structures, which were not detected in Cavin1 expressing cells (Fig. S6A-B). To address if this phenotype was due to defect membrane assembly, we purified the ΔDR1 (44-392) construct from mammalian cells and analyzed binding and insertion by QCM-D and monolayer adsorption experiments, respectively. We found that the ΔDR1 (44-392) appeared to bind slightly less to membranes (Fig. S6C), and that the kinetics of membrane insertion was significantly affected (Fig. 6D). However, at equilibrium the MIP was similar to full length Cavin (Fig. S6E). This showed that also in the context of the full-length protein, the DR1 region influence the kinetics of membrane insertion and that the observed phenotype on caveolae assembly in cells is likely due to impaired membrane assembly and insertion.

**Fig. 6.**
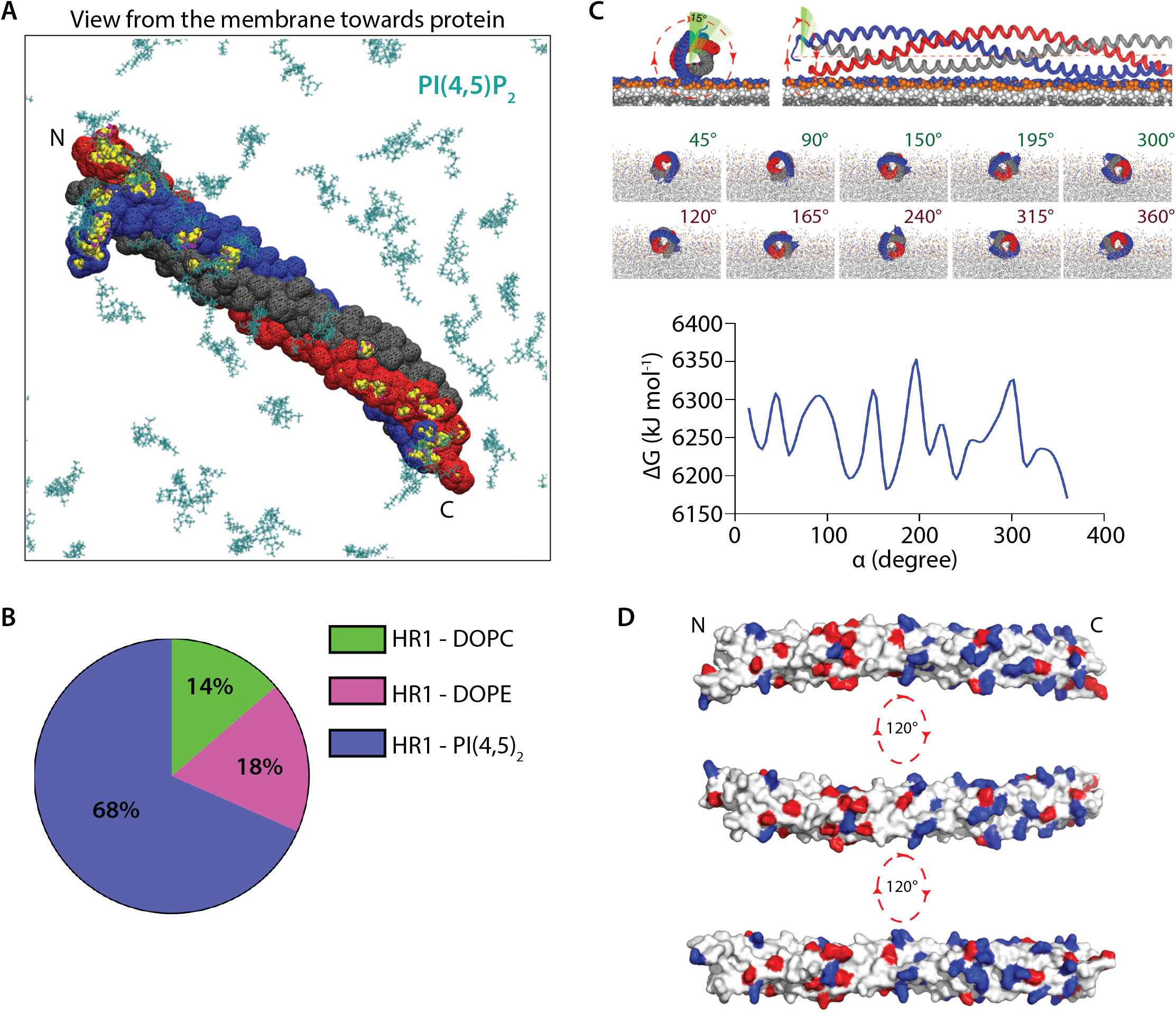
The HR1 domain binds via any of the three helices and orients parallel to the membrane interphase as revealed by MD analysis. **(A)** Representative bottom-up view from the membrane for all-atom simulations of HR1. Membranes consisted of DOPC, DOPE and PI(4,5)P_2_, but only PI(4,5)P_2_ (cyan) is shown for clarity. Residues highlighted in yellow were solvent-inaccessible and engaged in hydrogen bonding with the membrane, predominantly with PI(4,5)P_2_. **(B)** Pie chart showing the ratio of hydrogen bonding between HR1 domain and lipids present in the membrane based on all-atom MD simulations. The data was analyzed over the last 50 ns of 200 ns simulations. **(C)** All-atom simulations of HR1 rotation around its helical axis and associated free energy (ΔG) of protein-membrane interactions at different rotation angles. Lower panel in (C) shows the axial views at different angles, where maxima and minima of ΔG are color-coded in green and red, respectively, and correspond to maxima and minima energy states. **(D)** Electrostatic surface potential of the HR1 (red, negative charge; blue, positive charge).

### Molecular dynamics analysis reveals that membrane binding triggers partial helical separation of the HR1 and membrane insertion of the MIU

The mechanism for membrane insertion of the MIU in the structural context of the trimeric HR1 is difficult to envision. We therefore aimed to mechanistically dissect how the HR1 domain would bind and insert into membranes using molecular dynamics (MD) simulations. We limited the computational model to the HR1 domain, since the structure is described in detail, and it is the part of the protein known to interact with lipids. Using all-atom MD simulations, we found that the trimer instantly attached to the membrane surface consisting of DOPC:DOPE:PI(4,5)P_2_ (Fig. 6A). All three helices were engaged in hydrogen bonding once the trimer was horizontally attached and 68% of the bonds were formed between the trimer and PI(4,5)P_2_ (Fig. 6B). Especially, lysine residues in both the N- or C-terminal region of the HR1 domain contributed to most of the hydrogen-bonding interactions. This was confirmed by the solvent accessible surface area (SASA); a measure of whether residues in the protein are buried or solvent exposed (Fig. 6A). Next, we analyzed whether a particular orientation of the trimeric HR1 domain towards the membrane would be energetically favorable. For this, we simulated the rotation of the trimeric HR1 domain around its helical axis and evaluated the free energy of protein-membrane interactions by removing the protein from the simulation box at different rotation angles (Fig. 6C). This enabled us to compare at which orientation the protein was more stable, indicated by a lower energy state of the system. Thermodynamic integration revealed alteration of the free energy as the trimer underwent gradual rotation around its helical axis (Fig. 6C, lower panel). However, the difference between the lowest and the highest energy states was found to be of approximately 3% of the entire protein decoupling energy. This suggests that the HR1 domain will bind to the membrane in any of these most energetically favorable orientations although the precise helices engaged in binding can vary. In line with this, the positively charged residues interacting with PI(4,5)P_2_ are distributed homogeneously around the surface of the rod-like HR1 domain (Fig. 6D). This is in contrast to previously described membrane binding domains such as BAR domains, which have a preferred membrane interaction surface.

To address the temporal membrane binding of the HR1 domain, we performed coarse-grained MD (CG-MD) simulations. The HR1 domain was placed in a simulation box near lipid membranes composed of DOPC:DOPE, with or without PI(4,5)P_2_, over a time-frame of 2 µs. No binding was observed in the absence of PI(4,5)P_2_ (Fig. S7A), but in its presence, lysine residues in either the N- or C-terminus of the HR1 domain initiated binding to this lipid (Fig. 7A, left panel). Subsequently, interactions between PI(4,5)P_2_ and positively charged residues along the trimer surface resulted in a horizontal binding of the HR1 in relation to the membrane in all simulations (Fig. 7A, right panel). In this state, the HR1 was tightly packed towards the head group interphase. In agreement with thermodynamic integration data (Fig. 6C), the individual helices facing the membrane appeared to be different in between simulations (Fig. S7B). On average the distance between the lipid head groups and the center of the bilayer was 1.95 nm (26). The residues of HR1 closest to the membrane were on average 1.79 ± 0.15 nm from the bilayer center suggesting that the HR1 was shallowly buried in the membrane in between the head groups (Fig. 7B). This could account for the intermediate increase in surface pressure detected for the HR1 domain in the monolayer experiments (Fig. 5A). Interestingly, the CG-MD simulation also showed that the helices in the HR1 could become slightly uncoiled (Fig. S7B, simulation 6), suggesting that membrane interaction may induce partial separation of the helices in the HR1 trimer. When binding of single chain helices to the membrane was simulated using CG-MD (Fig. 7C), we observed that the residues of the individual helices were inserted into the membrane on average 1.42 ± 0.15 nm from the bilayer center (Fig. 7B). This suggested that partial uncoiling of the trimer would allow for the helices to insert deeper into the membrane (below the head group region) compared to the HR1 trimer. To elucidate if the binding and insertion of the HR1 involved separation of the coiled coil, we performed longer CG-MD simulations (36 µs). Interestingly, following the initial electrostatic interaction placing the HR1 horizontally towards the membrane, the individual helixes in the MIU started to separate (Fig. 7D). Moreover, this separation resulted in further insertion of the helixes into the membrane, 1.63 ± 0.25 nm from the bilayer center. (Fig. 7B and D), suggesting that membrane binding induces coiled-coil separation and insertion of the MIU of Cavin1. In agreement with this, we could observe a steep increase in the number of lipid interactions of the individual helices in the MIU and a concomitant decrease in the number of protein interactions (Fig. S7C and D). Taken together, we propose that HR1 initiates membrane binding of Cavin1 via electrostatic interactions with PI(4,5)P_2_, resulting in tight packing towards the bilayer. This enables unwinding of the coiled-coil and insertion of the MIU region in between the lipid head groups. Insertion is assisted by the repelling electrostatic nature of the DR1 and DR2 regions, resulting in the stable membrane insertion of Cavin1.

**Fig. 7.**
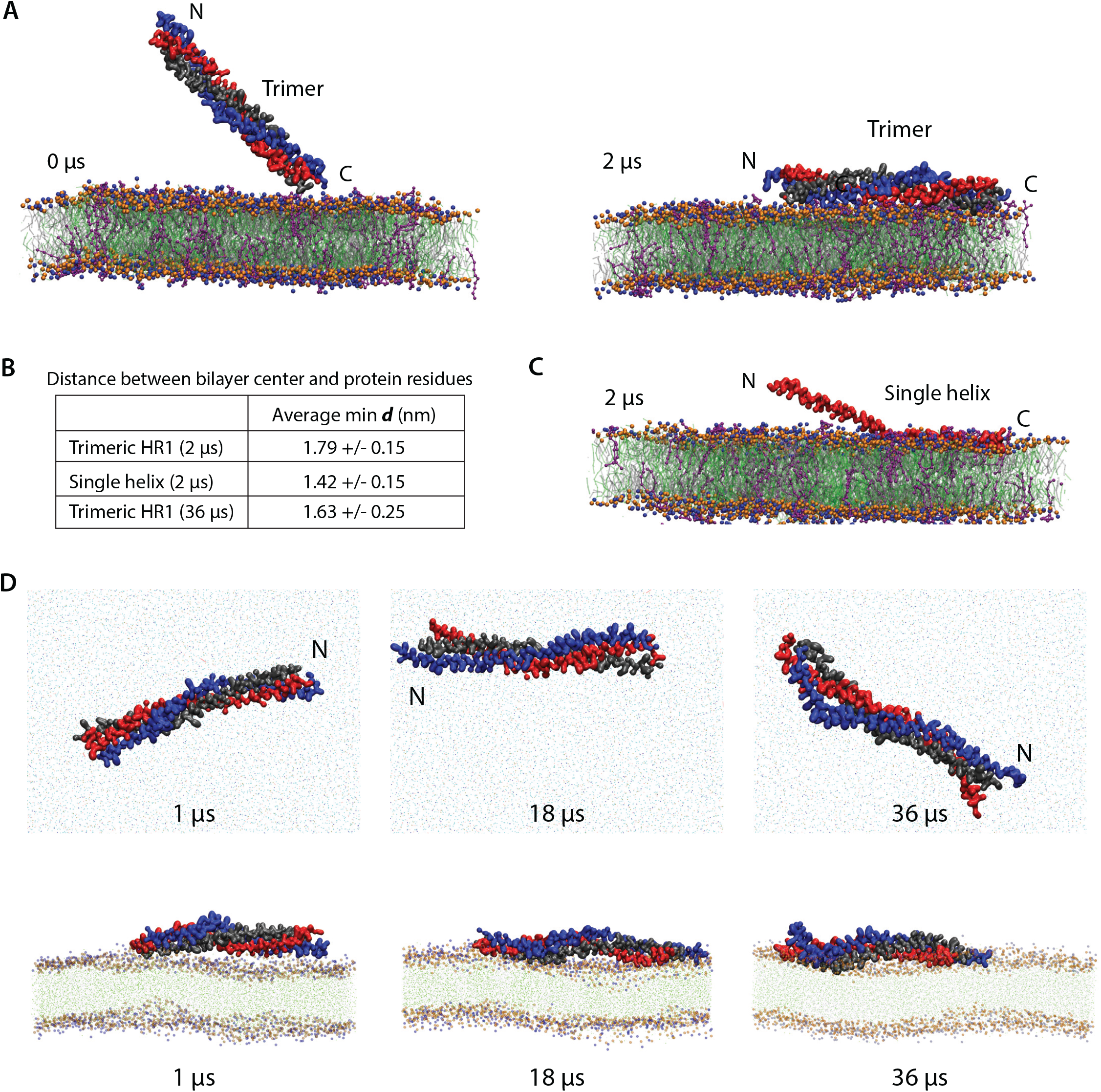
MD analysis shows that partial helix separation facilitate insertion of the MIU. **(A)** CG-MD simulations (2 µs) of HR1 membrane binding. Helices A, B and C are color-coded as blue, red and gray, respectively. Membranes consisted of DOPC (silver), DOPE (green) and PI(4,5)P_2_ (purple), where head group beads are shown in blue and orange. Binding was initiated by the N- or C-terminus (left panel) and HR1 was horizontally bound at the end of all simulations (right panel). **(B)** Overview of the distances between the bilayer center and the protein residues as determined in MD simulations over time and shown as average minimum distance (± SD). **(C)** Representative snapshot of CG-MD simulations (2 µs) as in (A) of monomeric HR1 domain (red) inserted into the membrane containing PI(4,5)P_2_ (purple). **(D)** Snapshots of the CG-MD simulations (36 µs) of HR1 membrane binding color-coded as in (A), where the upper and lower panels represent the top and side views of the system, respectively. The dots in both the upper and lower panel represent the beads of the membrane lipids color-coded as in (A).

## Discussion

The caveola-coat has eluded detailed architectural description for decades, but based on structural analysis of both caveolae and the components building up the coat, different models have been proposed (13, 14, 27). Validation and further improvements of these models rely on detailed structural understanding of the individual protein components at the membrane interface. In this work, we have studied the initial binding and assembly of Cavin1 at the membrane interface. We found that Cavin1 inserted into the membrane via a mechanism dependent on the DR1-HR1-DR2 interacting unit. Hereby, the HR1 undergo dynamic transition from a shielded coiled coil state in solution and a partially uncoiled state, where the MIU is partially buried in the membrane. Helix insertion has been shown to drive membrane curvature (28), but also to provide lipid specificity and targeting to specific cellular compartment as in the case of proteins containing amphipathic lipid packing sensor (ALPS) motifs (29). The described membrane insertion of Cavin1 would allow it to directly interact with lipid acyl chains. Such interactions could be linked to the recently reported lipid sorting activity of the caveolae coat, which was shown to be sensitive to the saturation level of acyl chains (6). The proposed mechanistic steps could provide specificity and regulation to the assembly process and enable Cavin1 to generate membrane curvature and interact with the integral part of CAV1 below the head group region of the membrane.

To characterize the properties of Cavin1 membrane binding in detail, we used purified full length and truncated versions of Cavin1 in combination with different membrane model systems. Whereby, we have been able to reconstitute and measure membrane binding and insertion of Cavin1 using a variety of biophysical techniques. The presented membrane insertion mechanism of Cavin1 was supported by the salt resistant binding of Cavin1 to SLBs as measured in real time using QCM-D. Furthermore, the IRRAS analysis showed that membrane binding of Cavin1 is coupled to decreased hydration of the lipid carbonyl groups, suggesting that Cavin1 inserted below the head group region of the membrane. Direct evidence for this comes from the surface pressure measurements using Langmuir lipid monolayers, which showed that Cavin1 rapidly inserts into lipid monolayers containing PI(4,5)P_2_. This was indicated by a steep increase in surface pressure, showing that the protein inserted in between the head groups of the lipids. The MIP of Cavin1 was higher than the monolayer-bilayer equivalence pressure, suggesting that Cavin1 spontaneously inserts into cellular membranes. The MIP is furthermore very similar to other proteins shown to insert into membranes using this methodology such as the BAR-domain of Bin1 (30) and Sar1p (31). Using analysis of truncated protein constructs we mapped the membrane interaction unit (MIU) to the C-terminal part of the HR1, which contains hydrophobic and positively charged residues key to membrane binding.

Using a combination of CG and all-atom MD simulations, we were able to dissect the binding interface of the HR1 and the hydrogen bonds formed with PI(4,5)P_2_. We found that binding was initiated and further supported by electrostatic interactions between PI(4,5)P_2_ and lysine residues along HR1 region resulting in tight horizontal docking of the HR1 domain. This is similar to previous simulations with a different lipid composition, POPC:POPS:PI(4,5)P_2_ (80:15:5 mol%) (6). Interestingly, we found that different helices of the trimeric HR1 were facing the membrane in the individual simulations. Careful analysis of the energies coupled to the orientation of the membrane bound HR1, revealed that the differences in ΔG following rotation of the HR1 were very small in comparison to the overall binding energy of the HR1. This is in agreement with the fact that the positively charged residues are distributed homogeneously around the surface of the rod-like HR1 trimer, showing that HR1 can bind in either of these orientations. This is dissimilar from other membrane binding domains, such as BAR domains, indicating that HR1-mediated membrane association of Cavin1 uses another type of mechanism.

Cavin1 has been proposed to bind as rod-like trimers based on the current structural knowledge. Yet, the striations detected on the caveolae bulb have been difficult to correlate to such Cavin1 rods, suggesting that the structure of Cavin1 might be flexible and change upon membrane association. Using CD-spectroscopy, we observed that membrane binding of Cavin1 induced structural rearrangements in the HR1, and IRRAS analysis revealed that the inclination angle of the membrane bound HR1 was slightly tilted. Furthermore, when CG-simulations of the HR1 region were performed for an extended time, we observed that the helices interacting with the membrane separated, which allowed the MIU to insert deeper in between the head groups. Indeed, the individual helices of the HR1 were able to insert deeper into the membrane as compared to the trimer. Taken together these results support the idea that the HR1 trimer might partially uncoil to expose hydrophobic residues, hidden inside the coiled core, during membrane binding and insertion. Helical uncoiling could be driven by the interactions between PI(4,5)P_2_ headgroups and the individual helices in a rotational movement competing with the interactions stabilizing the coiled coil. Similar drastic conformational changes have been observed in other proteins upon membrane binding. For example, membrane-driven exposure of amphipathic helices in small G-proteins (28) and the major helical rearrangement in pore forming proteins Bak and Bax upon membrane binding (32).

Our Langmuir trough data showed that membrane binding of the HR1 alone was inefficient in mediating membrane insertion. Instead, this was dependent on combining the HR1 with flanking disordered regions, suggesting that the interplay between the negatively charged DR1 and DR2 with the positively charged HR1 is important. Indeed, QCM-D data comparing combinations of HR1 with DR1 and/or DR2, showed that the softness of the membrane was dramatically altered in comparison to the full length Cavin1, unless all three regions were present. Indeed, deletion of the DR1 in the full-length protein affected the kinetics of membrane insertion and assembly of caveolae in cells, in agreement with previous reports (18). We propose that the DR1 and DR2 contribute to the destabilization of the HR1 coiled coil, thereby promoting insertion of the MIU. In line with this, the DR1 was shown to be required for Cavin1 induced membrane remodeling of spherical liposomes into membrane tubules. The disordered regions were proposed to influence assembly of Cavin1 via “fuzzy” electrostatic interactions with the helical regions. This induces a liquid-liquid phase separation and contributes to molecular crowding, which was proposed to drive membrane curvature and generate a meta-stable caveolae coat (18). Our *in vitro* and *in silico* data are consistent with this model and extend the implications of the weak electrostatic interactions between the DR1, DR2, and HR1 to membrane insertion. Taken together, the dynamic membrane insertion mechanism of Cavin1 described here provides a mechanistic basis for membrane-assisted regulation of the caveolae coat assembly.

## Materials and Methods

### Lipids

POPC (1-palmitoyl-2-oleoyl-*sn*-glycero-3-phosphocholine), PI(4,5)P_2_ (L-α-phosphatidylinositol-4,5-bisphosphate, porcine brain, ammonium salt), DOPC (1,2-dioleoyl-*sn*-glycero-3-phosphocholine), DOPE (1,2-dioleoyl-*sn*-glycero-3-phosphoethanolamine) and brain total lipid extract (FOLCH fraction, porcine) were purchased as lyophilized powder from Avanti Polar Lipids, Inc. (Alabaster, AL, USA).

### Protein purification

Cavin1 truncation proteins: 1-43, 44-155, 44-190, 1-155, 1-190, 1-100, 101-190 and 191-392, were purified as described previously (16). Proteins were expressed with N-terminal 6×His-tags in *E. coli* Rosetta pLysS or BL21(DE3)pLysS (growth in Terrific Broth media). Protein expression was induced with 1.0 mM IPTG at the exponential phase and incubated overnight at 20°C. TALON Superflow (Cytiva, Uppsala, Sweden) was used for affinity purification. Imidazole was removed by gel filtration chromatography using Sephacryl S-300 HR (Bio-Rad, Hercules, USA). Full length Cavin1 and ΔDR1 mutants were expressed and purified from suspension cells HEK-293F (Invitrogen, Carlsbad, CA, USA) as described previously (33). Cells were grown to 2– 3 × 10^6^ cells ml^-1^ on a shaker (160 rpm) at 37°C with 8% CO_2_ in 4 mM glutamine supplemented BalanCD medium (Irvine Scientific, Wicklow, Ireland). A total of 1 µg per one million cells of the plasmids containing CMV promoter and 3×FLAG-tagged geneswere mixed with a three-fold excess (w/w) of polyethylenimine MAX 40 kDa (Polysciences, Warrington, PA, USA) in 4 ml OptiPro (Invitrogen). The mixture was incubated for 20 min at room temperature before being added to the cell cultures. Cells were grown for two days with an addition of 5% BalanCD Feed (Irvine Scientific) per day. Cells were harvested, and lysed with 1% Nonidet P40 (Thermo Fisher Scientific, Waltham, MA, USA) for 15 min on ice. Following centrifugation at 20,000 *g* for 10 min, the supernatant was added to 3 ml anti-Flag (M2)-agarose (Sigma, St. Louis, MO, USA), and incubated at 4°C overnight. The gel matrix was transferred to a column and washed with 10 column volumes of 20 mM HEPES, 300 mM NaCl (pH 7.4). In order to remove Hsp70 chaperones, the matrix was incubated with a buffer containing 5 mM ATP, 20 mM MgCl_2_, 10 mM KCl and 0.1% Nonidet P40 for 2 h. The protein was eluted with 100 μg/ml of 3×FLAG peptide (Sigma). The eluted protein was adjusted to desired concentration via Vivaspin (Sartorius, Göttingen, Germany), analyzed by SDS-PAGE and snap frozen in liquid nitrogen, and stored at -80°C.

### Liposome co-sedimentation assay

The liposome co-sedimentation assay was performed as previously described (34). Briefly, FOLCH lipids or formulated lipid mixtures composed of DOPC:DOPE:PI(4,5)P_2_ (55:45:5 mol%) were dissolved to a final concentration of 1 mg/ml in chloroform: methanol (3:1 v/v). Lipids were dried under a stream of nitrogen, and rehydrated in 20 mM HEPES buffer, 150 mM NaCl (pH 7.4) followed by bath sonication (Transsonic T310, Elma Schmidbauer, Singen, Germany). Proteins were incubated with liposomes at a final concentration of 3 μM and 0.5 mg/ml, respectively, for 15 min at room temperature. The samples were centrifuged at 100,000 *g* for 20 min at room temperature. Then the supernatant and pellet were analyzed by Coomassie stained SDS-PAGE, and quantified using Image Lab software (Bio-Rad).

### SLB and QCM-D

Vesicles for foaming SLBs were prepared as described in liposome co-sedimentation assay. Except for the vesicles containing POPC:PI(4,5)P_2_ (95:5 mol%), which were extruded 11 times (Mini Extruder, Avanti) through a polycarbonate filter (Nuclepore Track-Etched Membranes, Whatman, Maidstone, UK) with 100 nm pore size. An AWSensors X4 unit (AWSensors, Valencia, Spain) equipped with a flow chamber was used to conduct the QCM-D measurements. Wrapped 14 mm (5 MHz, Cr/Au - SiO_2_, polished) sensors were used for all experiments. Each sensor was stored in 2% sodium dodecyl sulfate (SDS) overnight and treated with UV-ozone (Bioforce Nanosciences, USA) for 30 min prior to use. The frequency and dissipation changes for overtones 1, 3, 5, 7, 9, and 11 were all recorded, but only the third overtone was reported herein. POPC:PI(4,5)P_2_ vesicles (100 μl, 0.1 mg/ml) in 20 mM citrate, 50 mM KCl, 0.1 mM EDTA (pH 4.5) were injected in a continuous flow and SLB formation was monitored. After SLB formation, the chambers were rinsed with the buffer (20 mM HEPES, 150 mM NaCl, pH 7.4). After reaching a stable baseline, protein was injected into the chamber. Flow was paused once the protein solution had filled the sensor chamber and the system was allowed to reach equilibrium, before rinsing with the buffer. High salt treatment was done by rinsing the sensor surface with the same buffer but a higher salt concentration (300 mM NaCl) followed by rinsing with the initial buffer (150 mM NaCl).

### Lipid monolayer experiments

Lipid monolayer experiments were performed with either a custom-built round PTFE trough at Martin-Luther-Universität, Halle-Wittenberg (Ø 60 mm × 3 mm, Riegler and Kirstein GmbH, Potsdam, Germany) or a Microtrough G1 system at Umeå University (Ø 53 mm × 4 mm, Kibron, Helsinki, Finland). Both troughs were covered to prevent temperature and humidity loss and temperature of the subphase was controlled through a circulating water bath. Lipid mixtures consisting of DOPC:DOPE:PI(4,5)P_2_ (55:45:5 mol%) were prepared at total lipid concentration of 1 mM in chloroform:methanol (3:1 v/v). A microbalance equipped with a Wilhelmy plate was used to measure the surface pressure (π) and calibrated before each measurement. Lipid solutions were deposited onto the surface of the subphase (25 mM HEPES, 300 mM NaCl, 25 mM KOH, pH 7.4) to obtain the required initial surface pressure π_0_. The subphase was continuously stirred by a magnetic stirrer. After the solvent had been allowed to evaporate for 15 min and a stable monolayer was formed, the protein was injected under the lipid film directly into the subphase using a thin syringe needle (final concentration 50 nM). Curve analysis of Δπ/π_0_ plot provides synergy factor (*a*) as the slope of the linear regression +1. A positive *a* is indicative for attractive interactions between the lipid monolayer and the injected protein, while *a* = 0 would indicate a lack of interactions. The maximum insertion pressure (MIP) was determined from the Δπ/π_0_ plot through linear extrapolation to the x-axis, *i*.*e*., it corresponds to π_0_ at Δπ = 0 (22). The standard deviation of the MIP value was calculated according to the formula given in (22). Analysis of the adsorption curves was performed with Origin 8.1 (Origin Lab Corp., Northampton, MA, USA).

### Monolayer measurements with IRRAS

The Langmuir trough system used in combination with IRRAS (Riegler and Kirstein GmbH) included a circular sample (Ø 60 mm; 7.4 ml) and a rectangular reference trough (30 × 6 cm). The levels of the subphase (either H_2_O or D_2_O based) were controlled with an in-built laser and could be externally regulated via a pump system. The subphase was maintained at 20°C through a circulating water bath. The same procedure as above was followed for preparation of the lipid film. IRRAS experiments were conducted with a Bruker Vertex 70 FTIR spectrometer equipped with an A 511 reflection unit (Bruker, Karlsruhe, Germany) and an external mercury cadmium telluride (MCT) detector. The entire setup was enclosed and purged to keep the relative humidity constant. IRRA spectra of the films were acquired at various angles of incidence (between 25 and 70°) using parallel (p) and perpendicularly (s) polarized infrared light. 2000 scans were accumulated in p and 1000 in s polarization of the IR beam with a resolution of 8 cm^-1^ and a scanner frequency of 80 kHz. An additional zero filling factor of 2 was applied to the averaged interferograms prior to Fourier transformation. The single-beam reflectance spectra of the reference (R_0_) and the sample (R) trough surfaces were used to calculate the reflection absorption spectrum as lg (R/R_0_). Details for IRRA spectra simulation, band fitting parameters and principal component analysis are described in the supplementary materials.

### CD-spectroscopy

The secondary structure of Cavins was analyzed using a circular dichroism spectropolarimeter (JASCO J-810, Tokyo, Japan) at 25°C in the presence and absence of FOLCH liposomes. The final protein and liposome concentration was 3 μM and 0.5 mg/ml, respectively, in 25 mM HEPES, 150 mM NaCl (pH 7.4). A cuvette with a 0.1 cm path length was used to acquire the spectra, which were measured from 190–260 nm by averaging 8 scans of each sample at a bandwidth of 2 nm and a scan rate of 50 nm/min. All samples were incubated for 5 min until equilibrium temperatures is reached. The buffer and liposome only spectra were measured as the background signals that were subtracted from protein signal.

### Mass photometry

Mass and oligomerization measurements were performed on glass coverslips (No,1.5 H, 24×50 mm, Marienfeld) with a CultureWell™ reusable gasket (Grace Bio-Labs) placed on top of it and recorded on a mass photometer (Two^MP^, Refeyn Ltd, Oxford, UK). The gasket well was filled with the sample (25-100 nM) and data acquisition was performed using AcquireMP (Refeyn Ltd) for 60 s. Each measurement was repeated at least 3 times. The recorded videos were analyzed using DiscoverMP (Refeyn Ltd). The molecular weight was obtained by contrast comparison with Bovine serum albumin standard calibrants measured on the same day.

| Protein | Mass | $\sigma$ | Molecules count (%) | Oligomerization state |
| --- | --- | --- | --- | --- |
| **1-155** | 55 | 18.3 | 3306 (109%) | trimer |
| **1-190** | 32 | 8.6 | 481 (24%) | mono |
|  | 100 | 13.5 | 1268 (63%) | trimer |
| **45-155** | 48 | 13.2 | 1761 (96%) | trimer |
| **44-190** | 55 | 12.0 | 1692 (75%) | trimer |
|  | 157 | 10.9 | 129 (6%) | ennea |
| **1-100** | 60 | 12.7 | 2469 (98%) | trimer |
| **101-190** | 42 | 13.9 | 676 (55%) | trimer |

### Computational simulations

All simulations were performed using Gromacs 2018 software (35), and mouse Cavin1 HR1 domain structure (PDB: 4QKV) was used as the initial model (15). The membrane binding and insertion of HR1 domain, including trimer orientation and rotation, were predicated using coarse-grained molecular dynamics simulations (CG-MD). The hydrogen bonds, free energy changes and HR1 domain-membrane interactions were predicated by all-atom MD simulations. Details for software, scripts and parameters are described in the supplementary materials.

### Cell culture, transfection and live cell microscopy

PC-3 cells (ECACC 90112714) were maintained in RPMI medium (GIBCO, Thermo Fisher Scientific) supplemented with 10% fetal bovine serum and penicillin/streptomycin. For live cell, confocal imaging 400,000 cells were seeded on 1.5 high tolerance glass coverslips 25 mm (Warner Instruments, Hamden, CT, USA) 24 h prior to transfection. Caveolin1-RFP, Cavin1-GFP and Cavin1-ΔDR1-GFP were transiently transfected using Lipofectamine 3000 (Invitrogen) according to manufacturer’s instructions 16 to 24 h before the experiment. Live cell experiments were performed using a growth chamber (37°C, 5% CO_2_) in connection to a Nikon Eclipse Ti-E inverted microscope microscope (Nikon Instruments Inc. Tokyo, Japan), equipped with DU897 ANDOR EMCCD camera (Oxford Instruments, Abingdon, UK) and a Nikon CFI Plan Apochromat 60x Oil (N.A 1.40) DIC objective and a Nikon CFI Plan Apochromat 100x (NA 1.49). The TIRF objective was controlled by NIS Elements interface (Nikon Instruments Inc.). Images were prepared using Image J (36) and Photoshop CS6 (Adobe Inc, San Jose, CA, USA).

## Acknowledgments

We acknowledge the Biochemical Imaging Center and Umeå Centre for Electron Microscopy at Umeå University and the National Microscopy Infrastructure, NMI for providing assistance in microscopy. The computations/data handling were enabled by resources provided by the Swedish National Infrastructure for Computing (SNIC) at the Uppsala Multidisciplinary Center for Advanced Computational Science (UPPMAX), the Center for High Performance Computing (PDC), and the High-Performance Computing Center North (HPC2N) partially funded by the Swedish Research Council (grant agreement no. 2018-05973). This work was supported by the Kempe foundation, the Swedish Research Council, the Swedish Cancer Society, Wallenberg Centre for Molecular Medicine and the Medical Faculty at Umeå University as well as the European Research Council.

## Author contributions

Conceptualization: KCL, EL, MH, RL

Methodological conceptualization: HP, CS, AS, SH, AK, MH, RL

Investigation: KCL, HP, EL, SH, AK, JM, CS, VG, MH

Funding acquisition: MB, CSAB, RL

Supervision: MH, RL

Writing—original draft: KCL, RL

Writing—writing, review & editing: KCL, HP, EL, CS, MH, RL

## Data and materials availability

All data needed to evaluate the conclusions in the paper are present in the paper and/or the Supplementary Materials.

