## Supplementary Materials for "Membrane insertion mechanism of the caveolae coat protein Cavin1"

**This PDF file includes:**

Computational simulation programs, IRRAS analysis, scripts and related references

Figs. S1 to S7

#### Supplementary Materials and Methods

##### IRRA spectra simulation, band fitting parameters and principal component analysis

IRRA spectra of the protein containing monolayer recorded on a $D_2O$ subphase (Sigma) were simulated in the range of the amid I' vibration according to a three-layer model reported by Kuzmin et al.(1). Simulation and fitting of the multicomponent IRRA bands were performed as described in Schwieger et al. (2). The optical constant of the subphase $D_2O$ were taken from Berti et al. (3). The refractive index of the lipid/polymer film was set to 1.41 and its layer thickness to 2 nm. The band positions of the subcomponents were derived from second derivative spectra and revealed the contribution of three amide I' band components centered at 1662, 1642, and 1626 $cm^{-1}$. These components were assigned to one random and two α-helical secondary structure elements, respectively (4). According to literature, the lowest component could also be related to β-sheet amid I' vibrations (5). However, neither the X-ray structure nor CD spectra gave any indication for the presence of β-sheet components in the structure of Cavin1. Rather, we assigned the two lower spectral components to water exposed and water shielded faces of the helices, respectively, as reported for other coiled-coil helical arrangements (6). In analogy, we assumed that the amide bonds in helical structures facing the aqueous subphase are more water accessible and absorb at lower wavenumbers (1626 $cm^{-1}$), whereas the interaction with lipid monolayer reduces hydrogen bonds with hydration water leading to a shift to higher wavenumbers (1642 $cm^{-1}$). The band component assigned to amide I' vibrations of unordered structures (1662 $cm^{-1}$) was assumed to originate from isotropically distributed amid bonds and, therefore, simulated with an order parameter of $S = 0$ (corresponding to a tilt angle $\theta$ of 54° with respect to the interface normal). The polar angle between the helix main axis and the transition dipole moments of the helix amid I' vibrations was set to $\alpha = 38°$ (5). The main axis tilt angle of the two lower band components were set to be identical and varied in a least square Levenberg–Marquardt fit to determine the most probable helix orientation. Further fitting parameters were the full-widths at half-height (*fwhh*) of the three band components as well as the respective absorption coefficients $k_{max}$. The spectra were fitted in the range of 1610–1670 $cm^{-1}$and at angles of incidence, $\varphi$ = 30–70° in increments of 4°, where $\varphi$ = 49, 52, 55 and 58° were omitted from the fit, because of low reflectivity in the range of the Brewster angle. For better understanding and coherence with the MD simulation results, the determined tilt angles $\theta$, which are defined with respect to the interface normal, are translated into inclination angles $\gamma$, which are defined in relation to the plane of the interface ($\gamma$ = 90°-$\theta$). The confidence interval of the fit minimum was calculated as follows: the goodness of the fit was assessed as the sum of square deviations at each inclination angle γ (SSD). These values were weighted by their minimum with (N-p), were N = 360 is the number of fitted data points and p = 6 is the number of fitted parameters:

$$X = \frac{SSD - SSD_{min}}{SSD_{min}} - 1 \times (N - p)$$

The X values are F-distributed with 1 degree of freedom. In the given conditions all $X \leq 3.87$ (SSD $\leq 1.26 \cdot 10^{-5}$) are within the 95% confidence interval.

A set of IRRA spectra was subjected to principal component analysis (PCA) in order to identify subtle changes in the band shape correlated to hydration differences. The PCA was performed on vector-normalized spectra (to exclude contributions from intensity variations to the principal components) in the range of 1693–1770 $cm^{-1}$. All spectra were pretreated as follows: i) a reference spectrum of a bare $D_2O$ subphase was subtracted from all spectra, ii) a $D_2O$ vapor spectrum was subtracted in a way to best reduce the contribution of vibrational-rotational bands in the spectral region of interest, iii) a polynomial rubber band baseline was subtracted from the spectra (Software OPUS, Bruker, Germany). All analyzed spectra were recorded in s-polarization, at various angles of incidence, $\varphi$. Four sets of spectra were analyzed in a common PCA: i) and ii) spectra of a DOPC:DOPE and of a DOPC:DOPE:PI(4,5)$P_2$ monolayer before injection of protein (20 mN $m^{-1}$, s-polarization, $\varphi$ = 40°) and ii) and iv) spectra of the respective monolayers after injection of Cavin1 (1-190) (*ca.* 36 mN $m^{-1}$, s-polarization, $\varphi$ = 25–70°). The PCA was performed using the *princomp* function of MATLAB (Math-Works Inc., Natick, MA, USA).

**Computational simulations programs and scripts**

**Coarse-grained molecular dynamics simulations of unrestrained protein.** For the coarse-grained (CG) simulations, Martini 2.2 force field (7, 8) was used and the initial PDB structure ID

4QKV (9) was coarse-grained using the Charmm-GUI platform (10, 11). The CG lipid membrane was generated using the insane.py script (11). Each simulation consisted of 674 lipid molecules per leaflet. Two different membrane compositions with molar ratio of DOPC:DOPE:PI(4,5)$P_2$ (55:40:5) and DOPC:DOPE (60:40) was used in different simulations. The topology of different lipid molecules was obtained from the martini website (http://cgmartini.nl/). The initial simulation box dimensions were 21, 21, and 27 nm in the x, y, and z directions, respectively. Each simulated system was first energy minimized using the steepest descent algorithm. This was followed by four short equilibration runs (100,000 steps) with time steps of 1, 2, 5, and 10 fs, before the final production run. The final production run was performed using 15 fs time steps and the temperature was set at 37°C. A periodic boundary condition was applied for all simulations with semi-isotropic pressure coupling due to presence of lipid membrane in the system. An in-house python code was used to calculate the distance between membrane center and the nearest residue from the membrane center. The number of contacts between the membrane lipid molecules and protein helices was obtained using the 'gmx mindist' tool in Gromacs with a cut-off distance of 0.6 nm for each component. All MD simulation snapshot images were generated using the molecular visualization program VMD (12). Images representing the electrostatic surface potential of the HR1was generated using PyMOL.

**Rotational map of free energy changes via coarse-grained molecular dynamics simulations of restrained protein.** Free energy calculations were based on the restrained Cavin1 HR1 model. The protein molecule was placed in the vicinity of the membrane and the positions of its beads were fixed in all 3 dimensions with a 10,000 kJ/mole energy penalty. We repeated the simulation for 24 angles of the rotation applied to the protein, with a step of 15 degrees, in order to evaluate the difference in affinities of various Cavin1 sides to the membrane. The membrane itself was not restrained in any way and both the lipids could move within the bilayer, and the entire membrane was free to move further away or closer to the protein. The membrane was assembled with Packmol software (13) and the topologies and coordinate files (except the protein) were obtained from the martini website (8, 14). As it was crucial to avoid the self-consistency through the periodic boundary conditions, the size of the membrane was chosen to span over the box more than the doubled major length of the protein. Each leaflet of the membrane consisted of DOPC:DOPE:PI(4,5)$P_2$ (55:40:5). Water layer of 10 nm with neutralizing ions was added to the box, so that the inclined protein would not attach to the membrane with both ends through the periodic boundary conditions. After the equilibration stage (500 ns), where the timestep was gradually increased, sets of simulations (150 ns each) were performed in order to measure the free energy change associated with the removal of the protein from the simulation box, while it was in contact with the membrane. Thermodynamic integration was used with 20 intermediate steps – lambdas. At these steps van der Waals and electrostatic interactions of the protein with the rest of the system were gradually removed. That technique is well known in the field of computational chemistry, but in most cases applied to a small unrestrained molecule (15), either to modify it (*i.e.*, protonation state (16), or to measure solubility in several solvents (17)). Here, it was applied to measure the rotational free energy map, varying the contact spot at the interface of a protein and a membrane. Restraints of the protein were needed in order to reach convergence of the energy in a reasonable amount of simulation time. The gmx bar module of Gromacs was used in order to put the values from all lambdas together and calculate the free energy change between the two terminal states. The standard deviation of the free energies measured within each lambda was not statistically meaningful. The entire set of simulations for 24 angles of rotation included 504 simulations with a total time above 100,000 cpu-hours.

**Hydrogen bonds and protein-membrane interactions in all-atom molecular dynamics simulations**. All-atom simulations were performed in order to investigate the formation of the hydrogen bonds between the Cavin1 HR1 (unrestrained) and the membrane. The simulations were performed using CHARMM36 force field (18). The position of the protein, as well as the number of hydrogen bonds formed between the protein and the membrane (including more specific helices-membrane and lipids-protein bonds) were evaluated. The "gmx hbond" module was used to evaluate the number of hydrogen bonds formed between different groups of molecules. The all-atom simulations were equilibrated and followed by a production run lasting 100 ns. The latter was performed using 2 fs time step. Verlet cut-off scheme was used with the radii of 1.2 nm for both coulomb and van der Waals interactions. PME type of electrostatic interactions representation was applied. Force-switch modifier was also applied to the van der Waals interactions, with a radius of 1.0 nm. Nose-Hoover thermostat was used with a $\tau_t$=1.0. Parrinello-Rahman barostat with semi-isotropic coupling was used with the $\tau_p$=5.0, compressibility of $4.5\times 10^{-5}$ and reference pressure of 1 atmosphere. LINCS algorithm was applied to hydrogen bonds.

**Supplementary figure legends**

### Figure S1

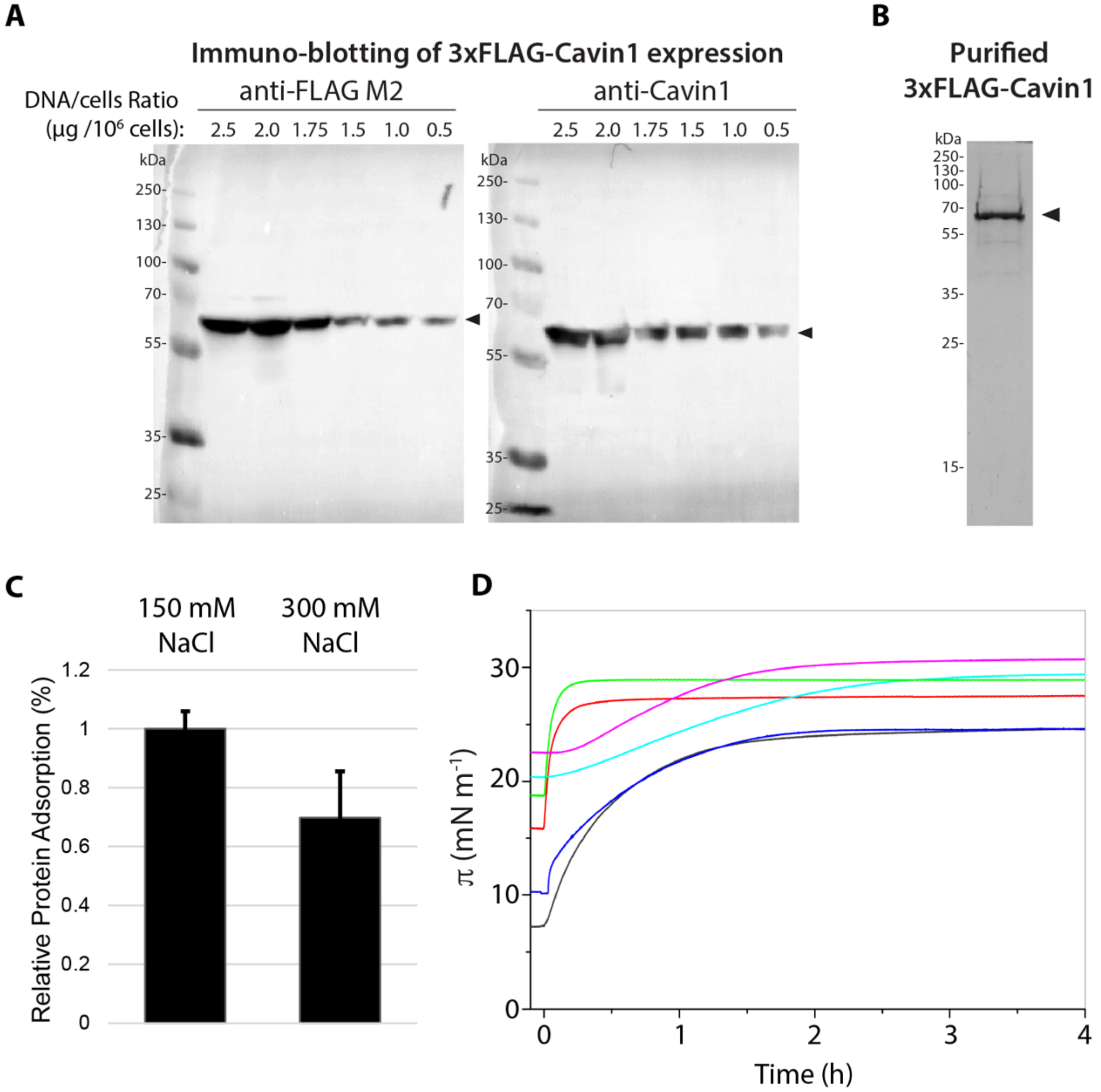


**Fig. S1. Purification of full-length Cavin1 expressed in HEK 293-F cells and monolayer adsorption experiments. (A)** Immunoblot of lysates from HEK 293-F cells transfected with different amount of 3×FLAG-Cavin1 expression plasmid as indicated. **(B)** Coomassie stained SDS-PAGE of purified 3×FLAG-Cavin1 protein. **(C)** Quantification of Cavin1 adsorption to SLBs as measured by QCM-D in buffer supplemented with 150 mM or 300mM NaCl. **(D)** Adsorption of Cavin1 to lipid monolayers. Lipid monolayers consisting of DOPC:DOPE:PI(4,5)$P_2$ (55:40:5 mol%) were prepared on a buffer subphase and Cavin1 was injected underneath the film at different initial surface pressure ($\pi_0$). The surface pressure ($\pi$) was recorded as a function of time.

This data was used to determine the maximum insertion pressure (MIP) of Cavin1 shown in Fig. 1F. **(D)** Quantification of Cavin1 adsorption to SLBs as measured by QCM-D in buffer supplemented with 150 mM or 300mM NaCl.

### Figure S2

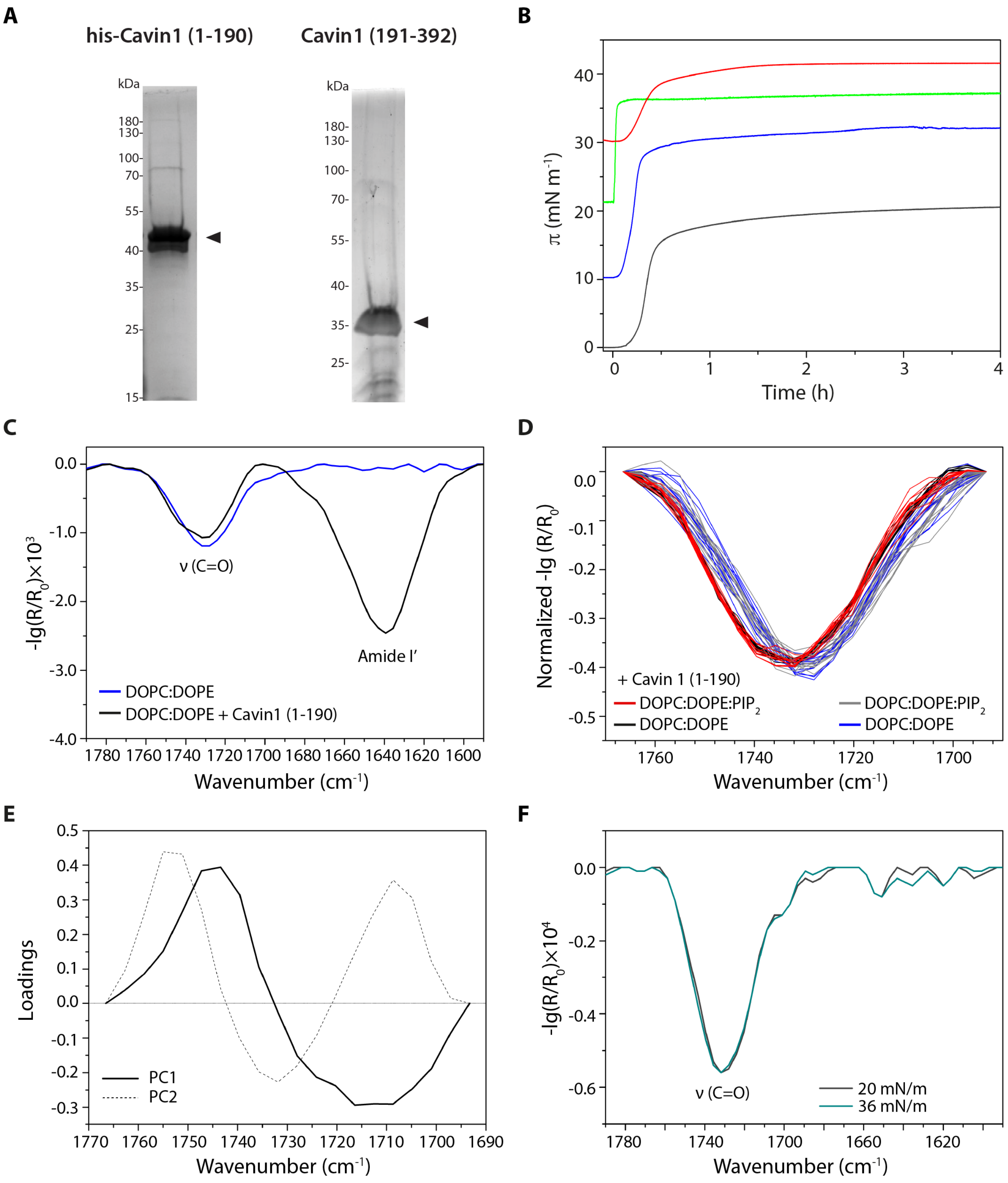


**Fig. S2. Purification, monolayer adsorption and IRRAS analysis of Cavin1 (1-190). (A)** Coomassie stained SDS-PAGE of purified Cavin1 (1-190) and (191-392) proteins from bacterial expression. (**B)** Adsorption of Cavin1 (1-190) to lipid monolayers. Lipid monolayers consisting of DOPC:DOPE:PI(4,5)$P_2$ (55:40:5 mol%) were prepared on a buffer subphase and Cavin1 (1-190) was injected underneath the film at different initial surface pressure ($\pi_0$). The surface pressure ($\pi$) was recorded as a function of time. This data was used to determine the maximum insertion pressure (MIP) of Cavin1 (1-190) shown in Fig. 2C. **(C)** IRRA spectra (1790–1590 $cm^{-1}$) of DOPC:DOPE (60:40 mol%) at an initial surface pressure of 20 mN $m^{-1}$. The characteristic C=O vibrational band at ~1730 $cm^{-1}$ originates from lipid ester groups. The amide I' band with a maximum at ~1640 $cm^{-1}$ indicates presence of Cavin1 (1-190) after its injection into the subphase. Spectra were acquired with p-polarized light at an angle of incidence of 40°. **(D)** Set of spectra used in the principal components analysis (PCA): vector normalized IRRA spectra in the C=O stretching vibrational region measured in s-polarization and at various angles of incidence of lipid monolayers prepared with and without PI(4,5)$P_2$ and in presence and absence of Cavin1 (1-190), respectively, as indicated. **(E)** Results of PCA of the spectra shown in (D): loadings (coefficients) of the first two principal components (PC1, solid line and PC2, dotted line), which represent 76% and 10% of the total variance of the data, respectively. PC1 is indicative for a shift of the band to lower wavenumbers and can be interpreted as an increase in hydration of the carbonyl groups. The scores of the individual spectra on PC1 and PC2 are shown in Fig. 2F. **(F)** IRRA spectra of pure lipid films of DOPC/DOPE/PI(4,5)$P_2$, without protein, at the surface pressure of 22 and 36 mN $m^1$. Spectra were acquired with p-polarized light at an angle of incidence of 40°.

### Figure S3

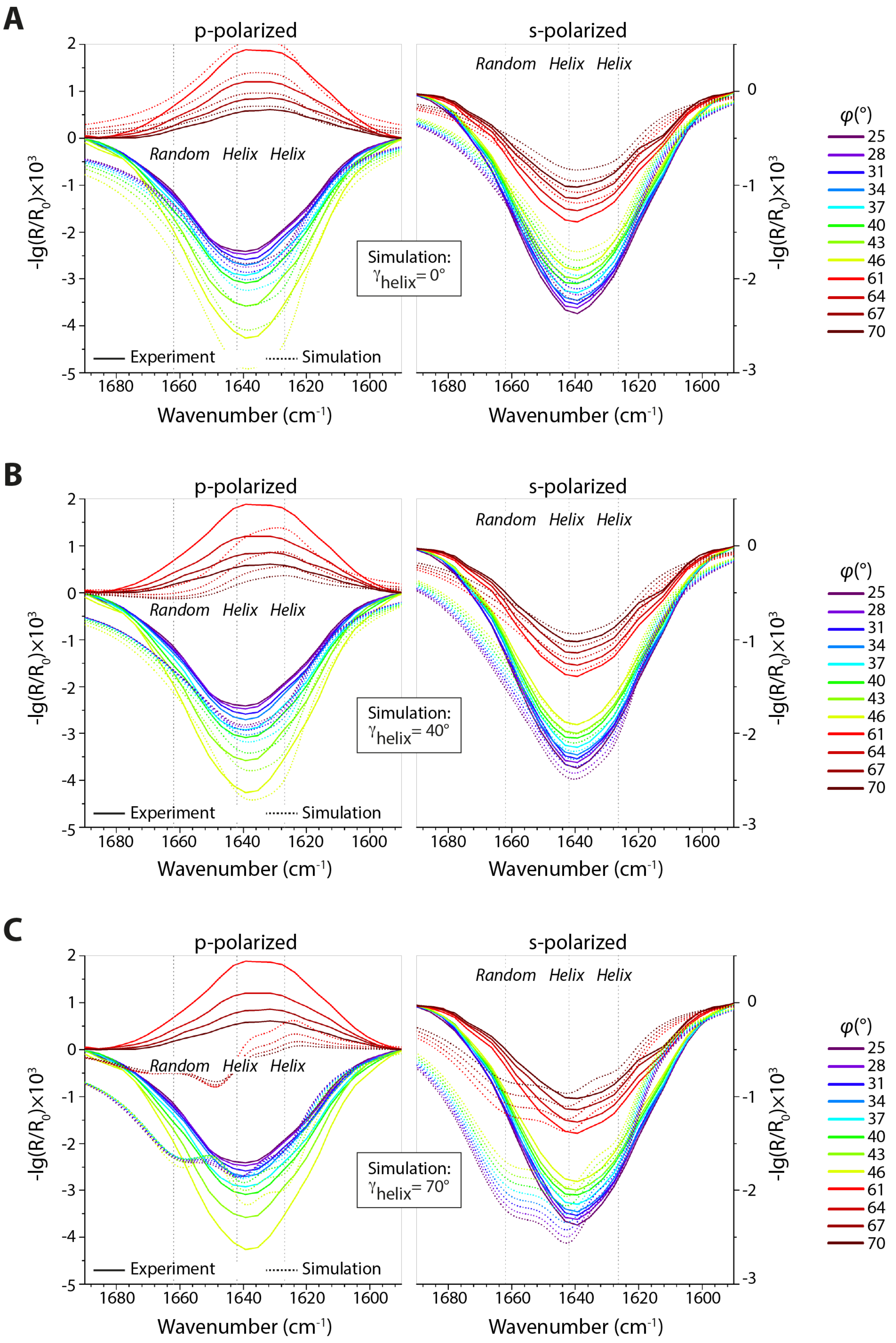
A
B
C
p-polarized
s-polarized
Random
Helix
Helix
-lg(R/R$_0$)×10$^3$
Wavenumber (cm$^{-1}$)
Experiment
Simulation
Simulation: $\gamma_{helix}$= 0°
Simulation: $\gamma_{helix}$= 40°
Simulation: $\gamma_{helix}$= 70°
$\varphi$(°)
25
28
31
34
37
40
43
46
61
64
67
70

**Fig. S3.** Evaluation of spectral fits of IRRA amid I' bands outside the 95% confidence interval of the minimum. Experimental (solid) and best fitting simulated (dotted) IRRA spectra of Cavin1 (1-190) adsorbed to a DOPC/DOPE/PI(4,5)$P_2$ monolayer, recorded at various angles of incidence $\varphi$ (see legend) and polarizations. The three band components used for simulation are indicated in the figure (vertical dotted lines). For SSD values of the shown fits see Fig. 3D. **(A)** The helix inclination angle was set to $\gamma = 0°$ which is identical to a completely parallel orientation of all helices to the lipid layer, which is only possible after HR1 uncoiling. Note that the shapes of the experimental IRRA amid I' bands could be reproduced by the fit but the intensities are not in perfect accordance. **(B)** The helix inclination angle was set to $\gamma = 40°$ (HR1 inclination: 40°). Note that the shape of the positive amid I' bands cannot be reproduced by the fit. **(C)** The HR1 inclination angle was set to $\gamma = 70°$, which is nearly upright (HR1 inclination: 74°). Note that that neither shape nor sign of the amid I' bands can be reproduced by the fit.

### Figure S4

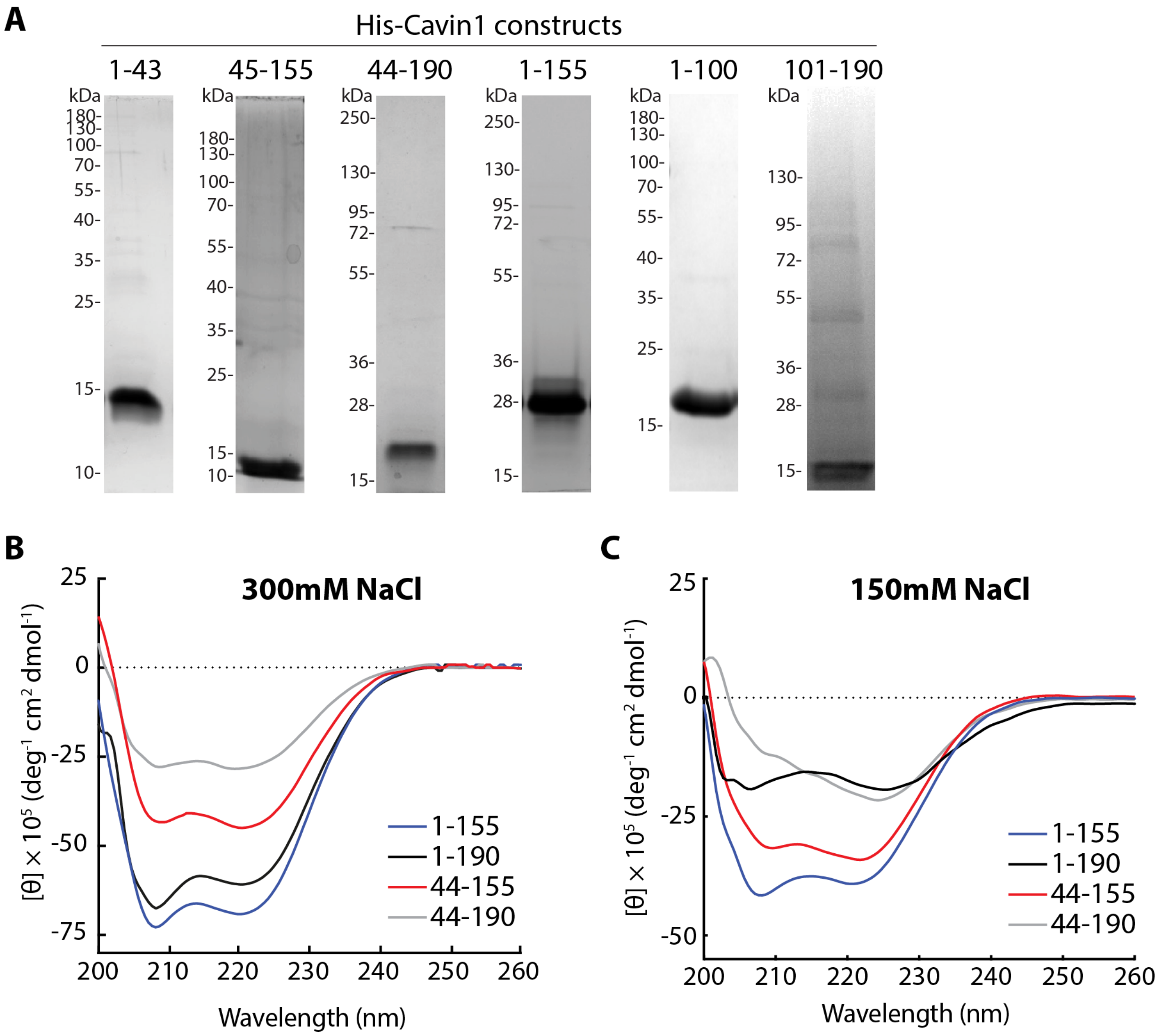


**Fig. S4. Purification and CD analysis of different truncated Cavin1 proteins. (A)** Coomassie stained SDS-PAGE gels of truncated Cavin1 proteins as indicated purified from bacterial: DR1, his-Cavin1 (1-44); HR1, his-Cavin1 (44-155); HR1+DR2, his-Cavin1 (44-190); DR1+HR1, his-Cavin1 (1-155), Cavin1 (1-100) and his-Cavin1 (101-190). **(B-C)** Far-UV CD spectra (195–255 nm) of Cavin1 constructs as indicated in 300 mM NaCl buffer **(B)** or 150 mM **(C)** NaCl buffer.

### Figure S5

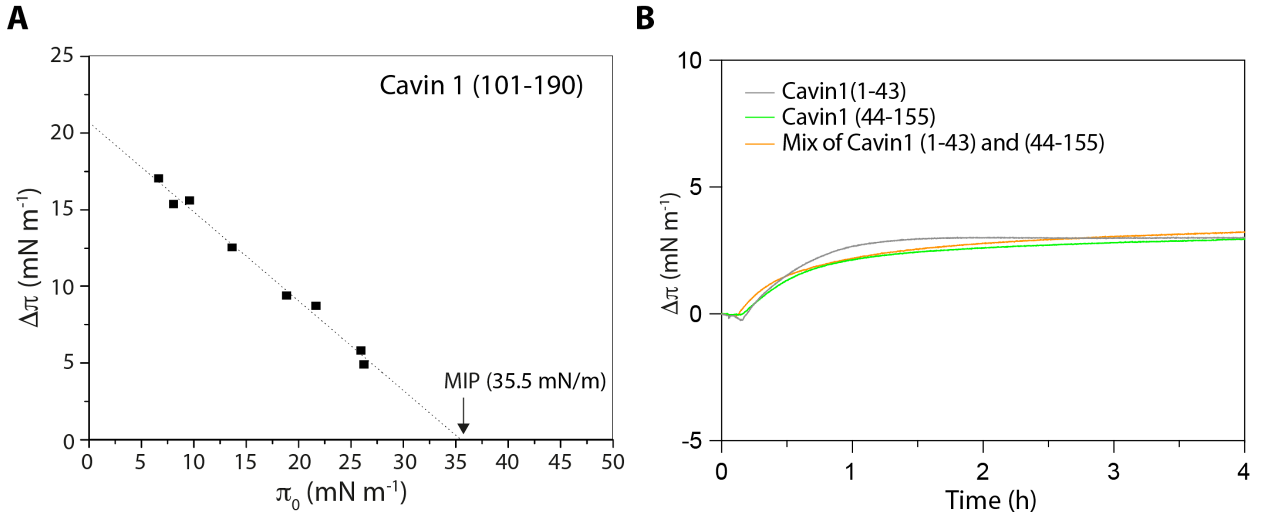


**Fig. S5. Monolayer adsorption of different truncated Cavin1 proteins. (A)** Cavin1 (101-190) adsorption to lipid monolayers was measured at different $\pi_0$. The MIP value was determined by extrapolation of the $\Delta\pi/\pi_0$ plot to the x-axis. **(B)** Adsorption of Cavin1 constructs to monolayer composed of DOPC:DOPE:PI(4,5)$P_2$ (55:40:5 mol%). Monolayers were prepared at $\pi_0$ = 20 mN $m^{-1}$. Following injection of the various truncates underneath the film, the surface pressure change ($\Delta\pi$) was recorded over time.

### Figure S6

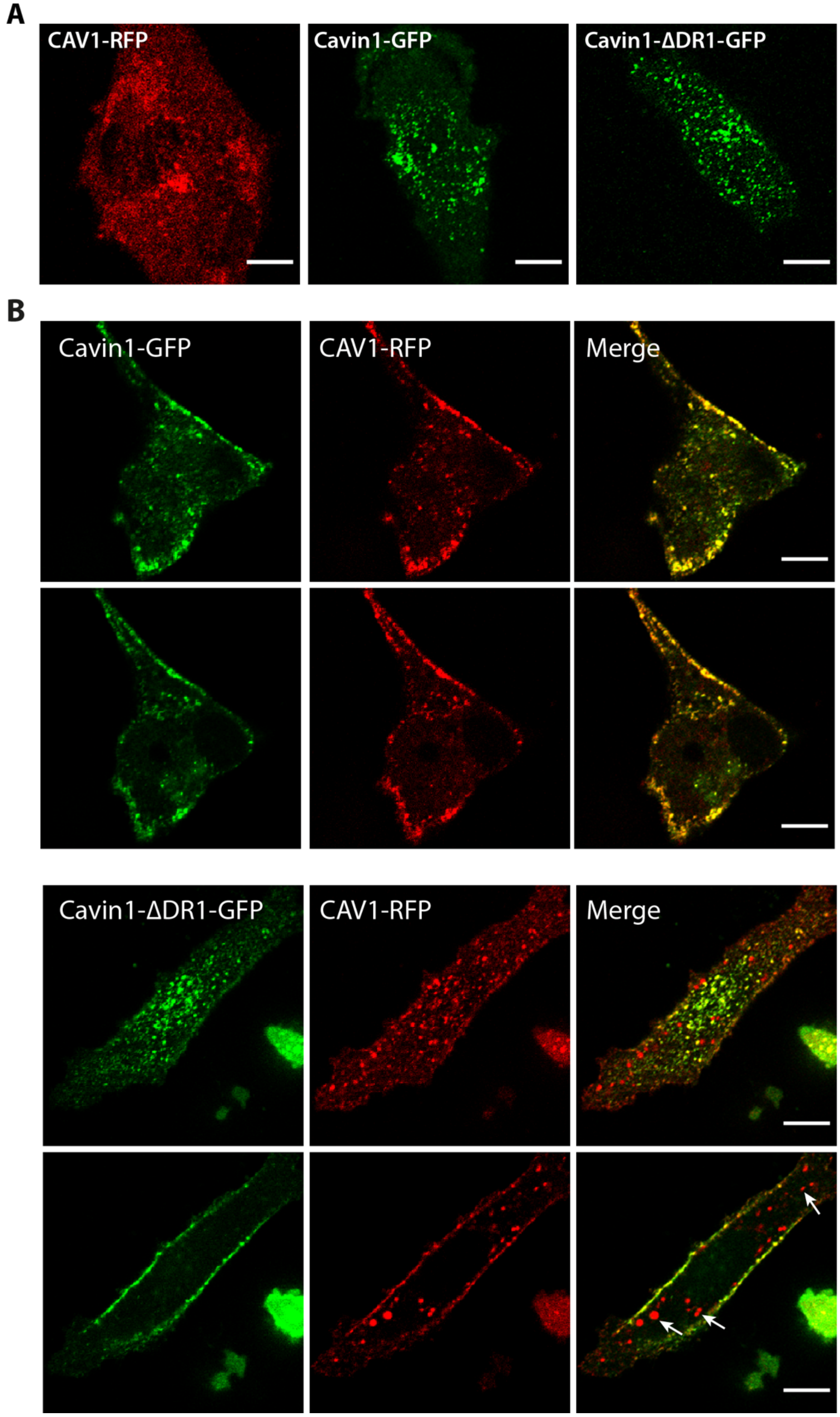
A
CAV1-RFP
Cavin1-GFP
Cavin1-ΔDR1-GFP
B
Cavin1-GFP
CAV1-RFP
Merge
Cavin1-ΔDR1-GFP
CAV1-RFP
Merge

### Figure S6

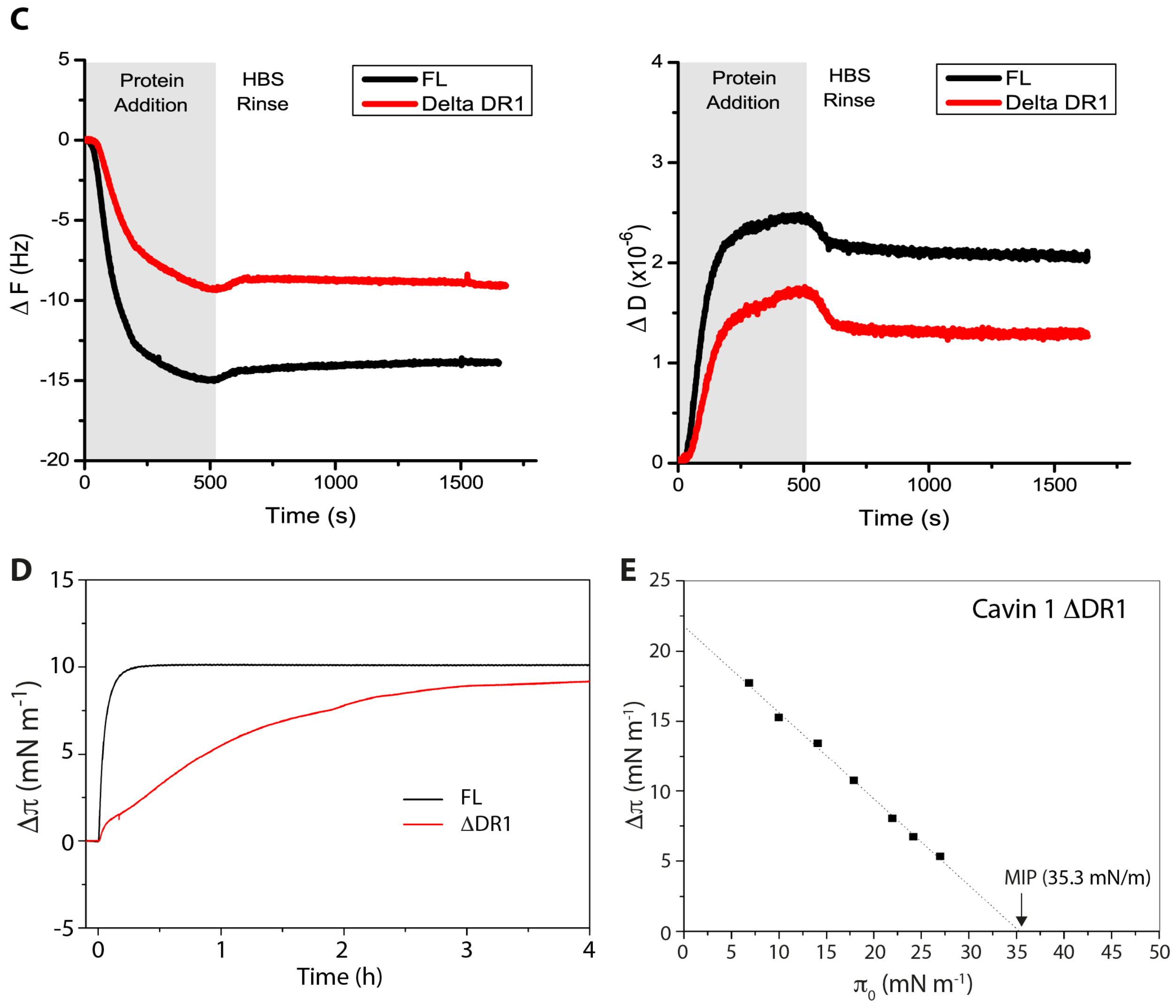


**Fig. S6. Deletion of the DR1 hampers membrane binding, and caveolae assembly in cells, but not membrane insertion. (A)** Representative confocal microscopy images of PC-3 cells expressing CAV1-RFP, Cavin1-GFP and Cavin1-ΔDR1-GFP as indicated. **(B)** Representative confocal microscopy images of PC-3 cells transfected with fluorescently tagged CAV1 (CAV1-RFP, red), full-length Cavin1 (Cavin1-GFP, green) or Cavin1-ΔDR1 (Cavin1-ΔDR1-GFP, green). Focal plane in the top panel shows protein localization at the basal membrane and in the bottom panel protein localization in the mid-section of a cell. White arrows highlight intracellular CAV1 structures devoid of Cavin1-ΔDR1-GFP. Scale bar =10 μm. **(C)** Frequency shift (ΔF) and Dissipation shift (ΔD) as a result of adsorption of ΔDR1 Cavin1 and full length Cavin1 onto SLBs measured by QCM-D. **(D)** Adsorption of ΔDR1 and full length Cavin1 to monolayer composed of

DOPC:DOPE:PI(4,5)$P_2$ (55:40:5 mol%). Monolayers were prepared at $\pi_0$ = 20 mN $m^{-1}$. Following injection of the various truncates underneath the film, the surface pressure change ($\Delta\pi$) was recorded over time. **(E)** Adsorption of ΔDR1 to lipid monolayers was measured at different $\pi_0$. The MIP value was determined by extrapolation of the $\Delta\pi/\pi_0$ plot to the x-axis.

### Figure S7

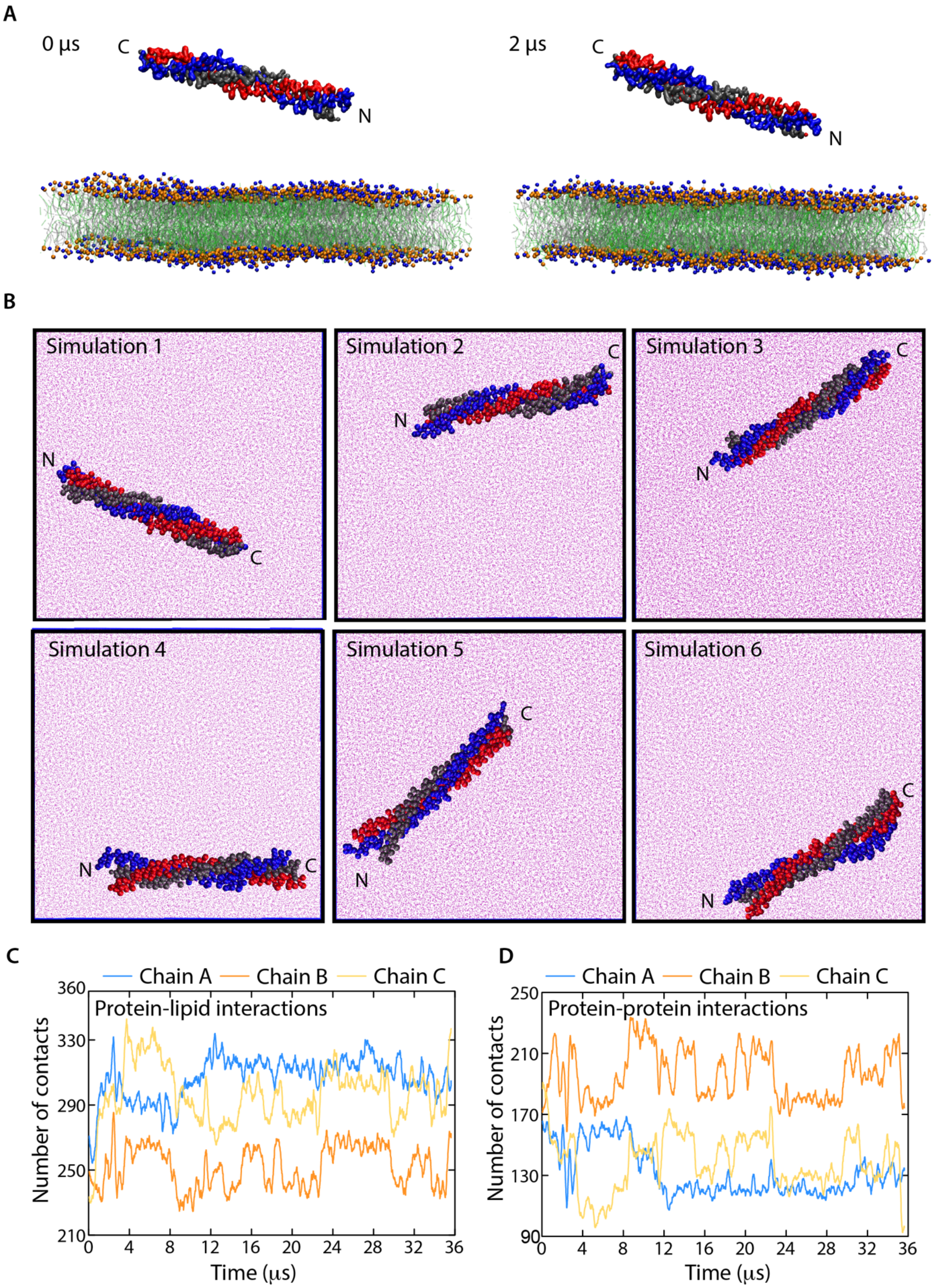
A
0 µs
C
N
2 µs
C
N
B
Simulation 1
N
C
Simulation 2
C
N
Simulation 3
C
N
Simulation 4
N
C
Simulation 5
C
N
Simulation 6
C
N
C
Chain A
Chain B
Chain C
Protein-lipid interactions
Number of contacts
360
330
290
250
210
0
4
8
12
16
20
24
28
32
36
Time (µs)
D
Chain A
Chain B
Chain C
Protein-protein interactions
Number of contacts
250
210
170
130
90
0
4
8
12
16
20
24
28
32
36
Time (µs)

**Fig. S7. Molecular dynamics simulations of the binding and orientation of HR1 domain at the membrane interphase. (A)** Coarse-grained molecular dynamics (CG-MD) simulations of HR1 domain binding to membranes consisting of DOPC (silver) and DOPE (green). Helices A, B and C of the HR1 crystal structure are color-coded as blue, red and gray, respectively. Lipid head group beads are shown in blue and orange. **(B)** Snapshots of bottom-up view from membrane toward protein-membrane interphase of trimeric HR1 domain in six CG-MD simulations. Membranes consisted of DOPC, DOPE and PI(4,5)$P_2$ (not shown). Helices A, B and C of the HR1 crystal structure are color-coded as blue, red and gray, respectively. **(C-D)** The number of contacts between the MIU (residues 101-155) of each chain of the protein with **(C)** the membrane lipids, and **(D)** the rest of the protein chains, during the 36 μs long CG-MD simulation. The pink dots represent water molecules.
